# Gene regulatory logic underlying the differentiation of mixed lineages in human thymic epithelial cells

**DOI:** 10.64898/2026.10.29.762771

**Authors:** Rishvanth K. Prabakar, Sarah R Chapin, Yong Lin, Stephen J. Staklinski, Deena M Netz, Madison E Lapine, Dominic Pruss, Todor Cvetanovic, Salome L Carcy, Joshua Torres, Alexander Grosh, Alexandra J Kiedrowski, Harvey I Pass, Ralph Mosca, Claire Regan, Jonathan Preall, Adam Siepel, Hannah V. Meyer

## Abstract

The thymus enacts central tolerance through thymic epithelial cells (TECs), which present self-antigens to developing T cells to remove self-reactive cells, preventing autoimmunity. Human TECs include populations that express lineage-committed programs corresponding to distinct peripheral tissues, yet how they acquire these programs and whether these TEC subpopulations are genetically related and conserved in mice remain unknown. Using matched single-cell gene-expression and chromatin-accessibility profiling of 126,687 human TECs, we identified 17 TEC subsets defined by their distinct expression profiles of 1,843 cell type restricted-antigens (CRAs) from 48 peripheral cell types. With a new phylogenetic reconstruction method, we show that lineage-committed TECs share a common ancestor, retain a thymus-specific mixed identity through co-expression of multiple CRA signatures, and are specified by coordinated activator and repressor regulons. We also identified a basement-membrane-collagen-rich state (BMC-TECs) with progenitor-like properties. Finally, mouse TECs shared the regulatory strategy but conserved few master transcription-factor drivers. This study mechanistically characterizes human TEC subpopulations and reveals the gene regulatory logic that leads to their distinct diversification and mixed identities in humans and mice.

## Introduction

As T cells develop in the thymus, strongly self-reactive cells are removed before they enter the circulation in a process called central tolerance, which is fundamental to immune homeostasis and protection from autoimmunity [1]. Thymic epithelial cells (TECs) provide diverse and specialized environments and broad representation of self-antigens that make this selection possible. However, the developmental origins and gene-regulatory mechanisms that generate human TEC diversity, including how TECs acquire full self-representation and whether these mechanisms are conserved across species, remain poorly understood.

Classical TEC subsets comprise cortical TECs (cTECs), which select developing T cells with functional T cell receptors and medullary TECs (mTECs), which present self-antigens and, together with co-stimulatory molecules, eliminate strongly self-reactive T cell clones. Within the medulla, expression of AIRE, a transcriptional regulator that induces expression of tissue-restricted antigens (TRA), and major histocompatibility complex (MHC) II distinguishes immature (mTECI; AIRE−, MHCII^lo^) from mature (mTECI; AIRE−, MHCII^hi^) subsets. Over the past two decades, central tolerance was attributed primarily to AIRE-expressing mTECIIs, whose stochastic expression of TRAs exposes developing thymocytes to peripheral self [1–4].

More recently, lineage-committed TEC subsets (CD24^hi^; *AIRE-*, MHCII^lo^), broadly classified as thymic mimetic cells, have been identified in mice and humans. Analyses of selected marker genes established their resemblance to peripheral cell types and provided a foundation for defining these specialized TEC states [5–9]. Examples include microfold- and corneo-like TECs, which express genes associated with intestinal M-cells and keratinization and cornified-envelope programs, respectively. In mice, lineage-committed TECs contribute to both tolerance and thymic functions beyond classical self-antigen presentation; for instance, tuft TECs contribute to invariant natural killer T cell development, whereas microfold TECs promote thymic B cell class switching [9–11]. Single-cell profiling and lineage tracing of mouse TECs have identified lineage-defining transcription factors (TFs) that establish these specialized gene expression programs [9–12]) and revealed distinct prenatal and postnatal lineage branches ([13]. Knockout of these TFs causes loss of the corresponding lineage-committed TEC [9–12] and can lead to autoimmune phenotypes in mice [10, 11]. However, whether these regulatory mechanisms and their consequences are conserved in humans remains unclear, particularly given species differences in subset abundance and lineage composition [8]. Addressing these questions requires a systematic, cell-type-resolved framework for defining and comparing peripheral programs in TECs.

Studies of human TECs have described five broad lineage-committed programs resembling muscle, neural, neuroendocrine, ionocyte and tuft cells [6–8]. However, their developmental origins and gene regulatory programs remain unresolved. It is unknown whether they follow shared or separate differentiation trajectories, and how chromatin and regulon architecture establishes peripheral cell signatures while preserving epithelial identity. Resolving human TEC gene regulation, lineage relationships, and cross-species conservation is therefore required to define how one epithelial compartment generates diverse cell states and how this diversity may contribute to human immune tolerance.

To address these questions, we systematically curated peripheral single-cell atlases to define cell type-restricted antigens (CRAs), establishing a cell-type-resolved framework for characterizing TEC identity. We then undertook single cell gene expression and chromatin accessibility profiling of human TECs from five pediatric donors and integrated these data with published thymic single-cell transcriptomics studies. Combining these resources resolved lineage-committed subsets, revealed thymus-specific co-expression of multiple peripheral cell signatures, and linked TEC subsets to regulon programs corresponding to their peripheral cell signatures. We also developed somatic mutation lineage tracing using these datasets, placing lineage-committed subsets on a shared differentiation trajectory and together with the CRA-guided annotation identified a progenitor-enriched BMC-TEC state. Lastly, cross-species analyses with mouse TECs identified a shared regulatory strategy but limited conservation of individual components. Together, these results show that human lineage-committed TECs arise from shared ancestry and retain a thymus-specific mixed peripheral identity under coordinated activator and repressor control, defining conserved gene-regulatory principles through which thymic epithelium diversifies to support immune tolerance.

## Results

### Cell-type-restricted antigen expression reveals 17 TEC subsets

We aimed to chart the full spectrum of TEC types in the human thymus and to understand the regulatory mechanisms that generate them. Doing so required two components: a census of cell types across human tissues and deep, single-cell measurements of both transcriptomes and chromatin accessibility in TECs.

To obtain the census of human cell types, we leveraged the CELLxGENE database [14] with over 62 million cells that represent 800 cell types and 370 tissues. We curated author-annotated labels into 48 broad, recurrent cell types across 45 human tissues by harmonizing labels, removing imprecise annotations, and requiring sufficient cells per tissue-cell-type category for reproducible pseudobulk estimates (Figure 1A, Supplementary Fig 1A, B; Supplementary Tables 1 and 2). Extending the concept of TRAs to individual cell types, we used the downsampled set of 1,558,164 cells to define CRAs and grouped them into cell-type-specific CRA modules with which to score TEC gene expression signatures (Supplementary Fig 1C, D). In total, we identified 1,843 CRAs (Supplementary Table 3), each specific to one of the 48 cell types (Figure 1B, marginal bar plot).

**Figure 1.**
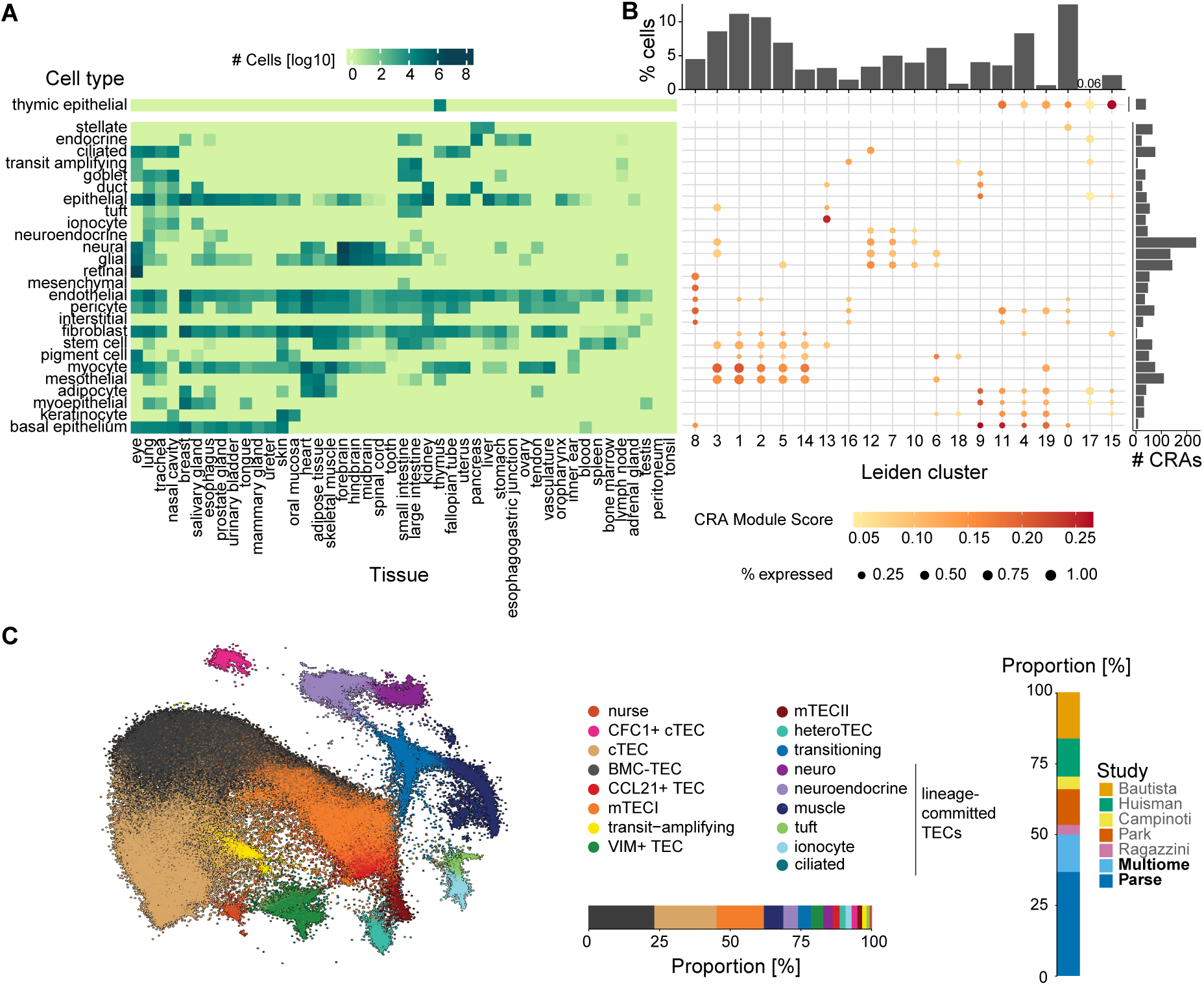
Cell-type-restricted antigen guided clustering of TECs. **(A)** Cell numbers of curated CELLxGENE cell types and tissues used for computing cell-type-restricted antigens (CRAs). **(B)** CRA module scores (dot color) and gene count (marginal bar chart, right) for integrated human TEC clusters (x-axis) and their associated cell counts (marginal bar chart, top); small proportions indicated by number. **(C)** single-cell ANnotation using Variational Inference (scANVI) embedding of integrated and CRA-annotated TEC types and cell type proportions and cell contributions per study (data contributed from this study in bold).

To resolve TEC-specific gene regulation at single-cell resolution, we profiled open chromatin (single-cell Assay for Transposase-Accessible Chromatin sequencing; scATAC-seq) and matched gene expression (single-nuclear RNA-sequencing; snRNA-seq) in flow-sorted EpCAM-positive TECs from five pediatric human donors (Supplementary Table 4) using the 10x multiome and Parse WT protocols; the matched scATAC-seq measurements are used for the lineage and regulatory analyses below (Supplementary Figs 2 to 5 and Supplementary Table 4). To increase power for rare TEC states, we reprocessed published human TEC scRNA-seq datasets [5–8, 15] (Supplementary Tables 5 to 7), using a unified pipeline, yielding batch-corrected and integrated gene counts across studies, donors and technologies (Supplementary Fig 6). Overall, this joint dataset comprised 126,687 TECs, with 63,067 (49.8%) contributed from this study (16,641 cells with paired snRNA-seq and scATAC-seq measurements and 46,426 Parse scRNA-seq cells).

Next, we computed CRA module scores across all TECs and integrated these with broad classical labels for TEC type (cTEC, immature and mature mTEC; Supplementary Table 8) in an iterative clustering workflow (*Methods: Single-cell RNA-seq analysis of TECs*). We retained 20 clusters (Leiden) at the coarsest resolution that distinguished 17 reproducible TEC states (Figure 1B, C; Supplementary Fig 6E-G).

CRA module scores identified two new TEC types: ciliated lineage-committed TECs and *vimentin*+ (*VIM*+) TECs with strong fibroblast and endothelial cell-specific CRA signatures, including high *VIM*, *IGFBP7*, *SPARC* and *SPARCL1* expression (Figure 1B, C; Supplementary Fig 6G). *SPARC*, *SPARCL1*, and *IGFBP7* regulate endothelial cell proliferation, angiogenesis, and vascular stability [16, 17], whereas *VIM* is an essential mesenchymal intermediate filament protein [18]. We also successfully recovered TECs expressing muscle, neural, neuroendocrine, ionocyte and tuft cell-specific CRAs, consistent with previous studies using individual marker genes (Figure 1B, C; [5, 6, 8]).

Two additional clusters expressed multiple CRA modules and could not be further resolved when clustering the full set of TECs (Figure 1B, C; Supplementary Fig 6E-G). One population co-expressed CRAs and marker genes characteristic of muscle, neuroendocrine, neural, and mTECI cell types, whereas the other combined epithelial programs spanning keratinocyte, basal, and goblet cell CRA signatures. We provisionally refer to these clusters as transitioning and heteroTECs, respectively, based on their mixed transcriptional profiles; because such coexpression may itself be thymus-specific, we retained these mixed states for follow-up rather than forcing them into single lineages.

Among classical TEC subsets, we found a transcriptionally distinct cTEC cluster with high expression of *CFC1*, *TP53AIP1* and *UBD* (*CFC1*+ TECs) and identified a transcriptionally distinct *CCL21*+ mTEC cluster, consistent with others [6] (Figure 1C). Separately, we identified a TEC cluster with a small but robust enrichment in the CRA stem cell module (Figure 1B), low expression of canonical cTEC, mTEC and lineage-committed TEC markers (*KRT15*, *CLU*, *AIRE*, *HLA-DR, CD24*), and high basement membrane-associated collagen expression including *COL4A5*, *COL4A6* and *COL8A1* (Supplementary Fig 6G). We refer to these cells as basement membrane collagen (BMC)-TECs (Figure 1C).

In summary, systematic curation of CELLxGENE enabled a CRA-guided survey of 48 peripheral cell-specific programs across human TECs, revealing 17 subsets that reflect the total diversity of TECs in humans. We recovered established classical and lineage-committed populations, identified three new TEC states—ciliated, VIM+ and BMC-TECs—and resolved transitioning TECs and heteroTECs as distinct mixed-lineage states.

### Human lineage-committed TECs share a common ancestor

We next asked whether lineage-committed and classical TECs arise through separate developmental routes or share ancestry, which would distinguish independent acquisition of peripheral cell signatures from diversification of a common thymic epithelial lineage. In mouse models, genetic reporters have established TEC progenitor–progeny relationships across independent studies (Supplementary Fig 7A; [11, 13, 19]), demonstrating that tuft and muscle lineage-specific TECs can diverge before the late *AIRE-expressing* mTECII stage, whereas mouse microfold and corneo lineage-specific TECs arise from progenitors that have expressed Aire. Because analogous genetic reporters cannot be used in primary human tissue, we developed a new phylogenetic reconstruction method that uses naturally occurring somatic variants in scATAC-seq reads as endogenous lineage markers and assigns cell identities from paired snRNA-seq profiles (Figure 2A). The method pseudo-bulks scATAC-seq reads by matched transcriptomic cell type, calls variants followed by stringent filtering to distinguish somatic from germline variants, and uses Bayesian and maximum-likelihood inference to estimate relationships among TEC subsets.

**Figure 2.**
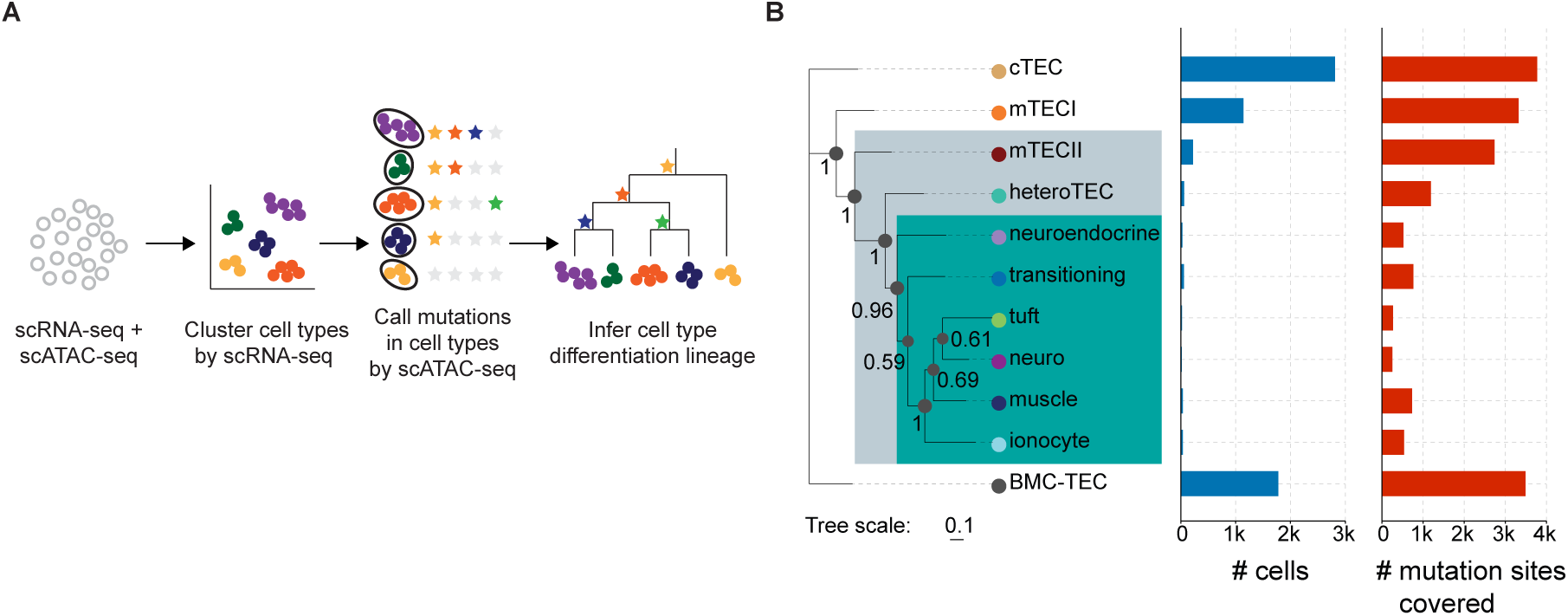
Human lineage-committed TECs share a common progenitor. **(A)** Methodology for inferring cell lineages from paired snRNA-seq and scATAC-seq data. **(B)** Rooted phylogeny of human TEC subsets (from 3,477 TECs), with the number of cells and callable mutation sites per cell type. Shared clades of lineage-committed TECs (teal) and mTECII common ancestor (grey) are highlighted. Numbers on branch points indicate posterior probabilities.

The previous mouse TEC snRNA-seq and scATAC-seq multiome dataset provided an ideal benchmark for our new method [19]. Its high per-base sequencing coverage (mean 2.44x/bp; Supplementary Fig 7B) yielded sufficient, high-confidence somatic variant calls per transcriptome-defined cell type to reconstruct a topology that could be compared with reporter-derived lineage tracing (Supplementary Fig 7A; [11, 13, 19]). Our phylogenetic reconstruction of mouse TECs placed tuft, ciliated and enterocyte TECs before the late AIRE-expressing mTECII stage, whereas other lineage-committed TECs clustered closer to this stage (Supplementary Fig 7C). Recovery of this established pre-mTECII placement of tuft TECs [10, 12] and the reported prenatal versus postnatal lineage splits [13] gave us confidence to apply our approach to human TECs.

Two of our three human donor datasets with paired snRNA-seq and scATAC-seq profiles passed the quality criterion observed for the mouse TEC dataset, with mean per-bp read coverage of 8.8 and 4.4 (Supplementary Fig 8A, B). We pooled the scATAC-seq reads aggregated by cell type and sample to call high-confidence somatic variants (Supplementary Fig 8C-K) and reconstruct cell-type phylogenies using Bayesian and maximum-likelihood inference. These analyses first yielded unbiased, unrooted topologies (Supplementary Fig 9A, B). We rooted the trees on cTECs, which form a distinct epithelial lineage from mTECs. The resulting phylogeny placed mTECI and mTECII along the classical medullary branch in an order that is consistent with the experimentally supported immature-to-mature transition, while the lineage-committed clade shared deeper ancestry with mTECII (Figure 2B, Supplementary Fig 9C). Transitioning and neuroendocrine lineage-committed TECs were closely related, as were neural, tuft, and ionocyte lineage-committed TECs. BMC-TECs formed a distinct branch in the phylogeny, raising the possibility that they represent a progenitor state, which we investigate further below. Maximum-likelihood reconstructions supported the same topology (Supplementary Fig 9D, E). The number of cells and callable mutation sites varied across cell types (Figure 2B), which reduced support at some internal nodes, but major lineage-level relationships were consistent across methods and samples.

Thus, in primary human thymus tissue, somatic variant-guided phylogenetic analysis of paired scATAC-seq and snRNA-seq data revealed that lineage-committed TECs form a shared clade with a common ancestor and share deeper ancestry with classical mTECII, supporting diversification from a shared thymic epithelial lineage.

### Human lineage-committed TECs acquire CRA expression on a common background

The shared ancestry of lineage-committed TECs raised the possibility that they retain a common gene expression background while acquiring distinct peripheral cell signatures. We therefore compared CRA expression across lineage-committed TECs, classical TEC subsets (cTEC, mTECI, and mTECII), and peripheral cell types.

Across the expression thresholds examined, mTECIIs expressed the greatest number of protein-coding (PC) and CRA genes of any TEC type, consistent with AIRE-driven promiscuous gene expression (Figure 3A). Lineage-committed TECs expressed fewer of these genes than mTECIIs but more than mTECIs and cTECs. By contrast, the expression of housekeeping genes was consistent across TEC types (Supplementary Fig 10A). These patterns replicated in each individual dataset (Supplementary Fig 10B-D). Relative to mTECII promiscuity, lineage-committed TECs adopted differentiated states dominated by CRAs corresponding to specific peripheral cell types. The PC and CRA genes expressed in lineage-committed TECs but not in mTECIs and cTECs were therefore consistent with a lineage-committed program superimposed on classical TEC states.

**Figure 3.**
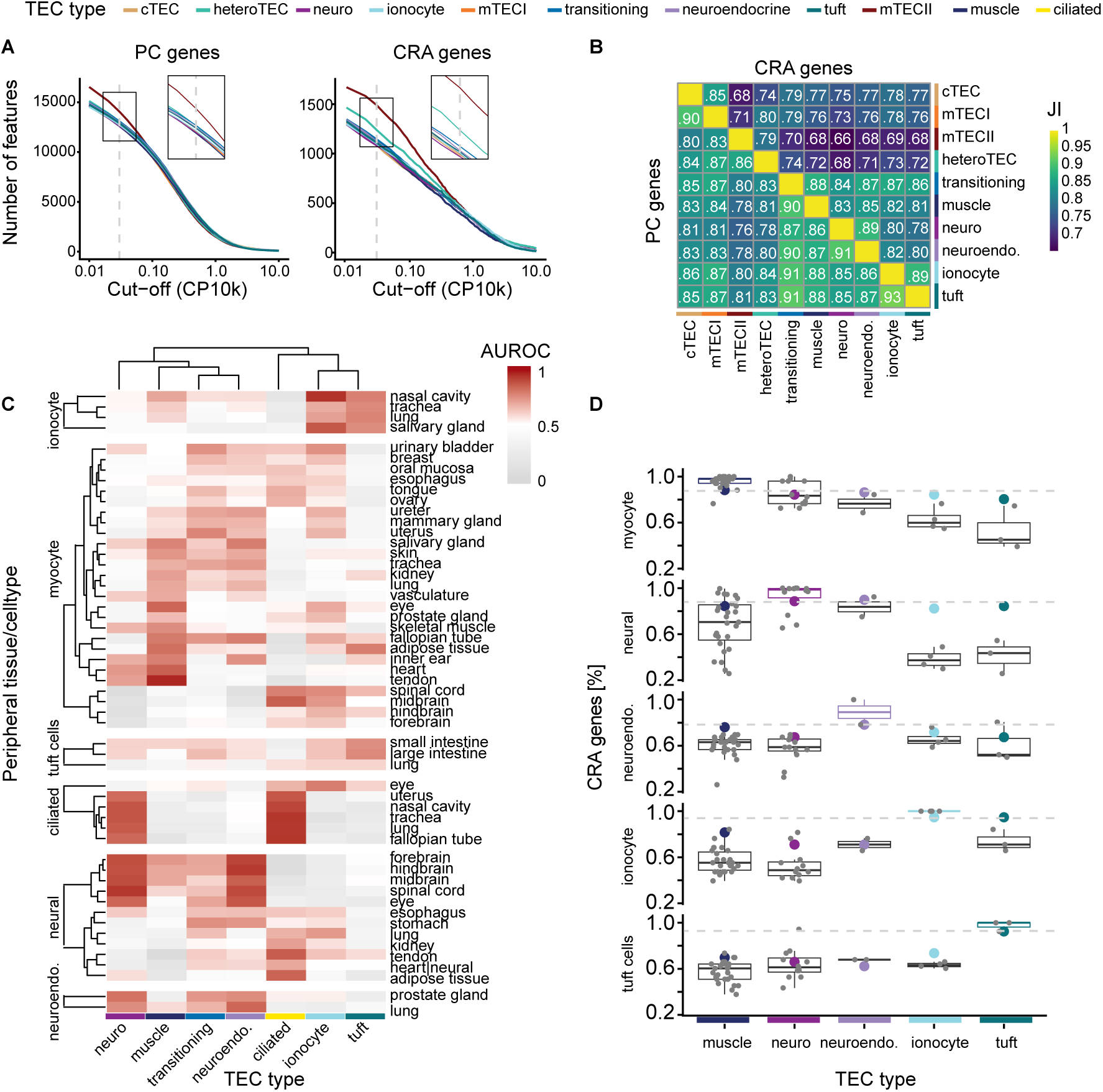
CRAs are shared between lineage-committed TECs. **(A)** Number of expressed protein coding (PC) and CRA genes across expression thresholds. Gray line indicates a threshold of 0.03, a representative value used for all downstream analyses. **(B)** Jaccard indices (JI) quantifying the number of shared PC and CRA genes between TEC types. **(C)** Area under the receiver operating characteristic curve from MetaNeighbor neighbor-voting analyses comparing lineage-committed TECs with their peripheral cell-type counterparts. **(D)** Fraction of CRA genes expressed in lineage-committed TECs from the integrated human TEC dataset (bold, colored dots) and peripheral cell types from the CELLxGENE dataset (box and gray scatter plots). The values conveyed by the dots and boxplots indicate the fraction of CRA genes corresponding to each of the peripheral cell type rows specified on the y-axis that are expressed by each cell-type-tissue combination. Peripheral tissue counts for each cell type: muscle: *n* = 27, neuro: *n* = 12, neuroendo. (neuroendocrine): *n* = 2, ionocyte: *n* = 4, tuft: *n* = 3.

We next characterized this program quantitatively. Using a representative expression cutoff of 0.03 (gray line in Figure 3A), we counted expressed genes in pseudo-bulk transcriptomes for each TEC type. Because binarized calls can be sensitive to cluster purity, we performed a computational mixing experiment (*Methods: Robustness of expressed gene counts to cluster annotations* Supplementary Fig 11) which confirmed that expressed-gene counts were robust to analytical and parameter choices and not inflated by mis-assignment. We then measured sharing of PC and CRA sets between TEC types using Jaccard indices (Figure 3B). As a control, housekeeping-gene sharing was uniform (empirical p value = 0.224; Supplementary Fig 12); by contrast, PC and CRA overlap was highest within lineage-committed TECs and exceeded pairwise comparisons involving only classical subsets (empirical p values = 0.162 and 0.048 for PC and CRA, respectively). This pattern supports a shared lineage-committed program rather than independent one-to-one peripheral mimicry in each TEC subtype.

To quantify the transcriptomic similarity of lineage-committed TECs to their corresponding peripheral cell types, we applied a neighbor-voting meta-analysis approach [20, 21] using the peripheral cell-type CRA modules (defined in Figure 1B). While each lineage-committed TEC was most similar to its respective peripheral cell type (Figure 3C Supplementary Fig 13), we also detected cross-lineage similarity; for example, muscle TECs also shared similarity to peripheral neural, neuroendocrine, and ionocyte cells, while ionocyte TECs shared similarity to peripheral neural, ciliated, and muscle cells. These patterns suggest that lineage-committed TECs share CRA modules associated with multiple peripheral cell types.

We therefore asked whether co-expression of multiple CRA modules was specific to the thymus. Quantifying CRA fractions in lineage-committed TECs versus peripheral cells from the curated CELLxGENE dataset (Figure 3D, Supplementary Figs 14 and 15), we found that each lineage-committed TEC expressed a smaller fraction of the CRAs corresponding to their peripheral cell type (in-group CRAs) but, surprisingly, also CRAs corresponding to other peripheral cell types (out-group CRAs), resulting in a broadened CRA profile that departs from a strict peripheral cell-specific match. For instance, muscle TECs expressed only 88.2% of myocyte CRAs, while peripheral muscle cells, on average, expressed 95.9% ± 4.93% of myocyte CRAs. Conversely, muscle TECs expressed large fractions of ionocyte (81.6%), neuroendocrine (76.1%), neural (84.4%), and tuft (69.8%) CRAs, whereas peripheral muscle cells averaged lower fractions for the same sets (*μ* =56.9% ± *σ*=10.3%, 60.5% ±8.84%, 68.5% ±22.1%, and 57.6% ±9.25%, respectively).

Together, these results define a thymus-specific expression pattern in which lineage-committed TECs retain CRAs associated with multiple peripheral cell types. Considered alongside their shared phylogenetic ancestry, this crosslineage CRA expression is consistent with retention of a common transcriptional background during diversification, rather than independently arising lineage-committed gene regulatory programs.

### TECs express peripheral cell regulons to establish lineage commitment

We next explored the gene regulatory framework that led to shared gene and CRA expression amongst lineage-committed TECs using our paired scATAC-seq and snRNA-seq data. This enabled us to relate chromatin accessibility to the gene expression programs that specify lineage-committed TECs and to test whether commitment reflects coordinated regulatory circuitry rather than the expression of select peripheral cell type-specific marker genes.

We first summarized chromatin “openness” by quantifying the fraction of fragments aligned to major genomic compartments (Figure 4A, Supplementary Fig 16). HeteroTECs and mTECIIs showed the largest fraction of open fragments in intergenic regions and relatively fewer open fragments in untranslated regions (UTRs) and transcription start sites (TSSs), matching the elevated epigenetic noise reported for mature TECs [19]. By contrast, lineage-committed TECs more closely resembled immature mTECI cells, with a larger share of open fragments in gene-proximal regions (TSS, UTR, coding sequence).

**Figure 4.**
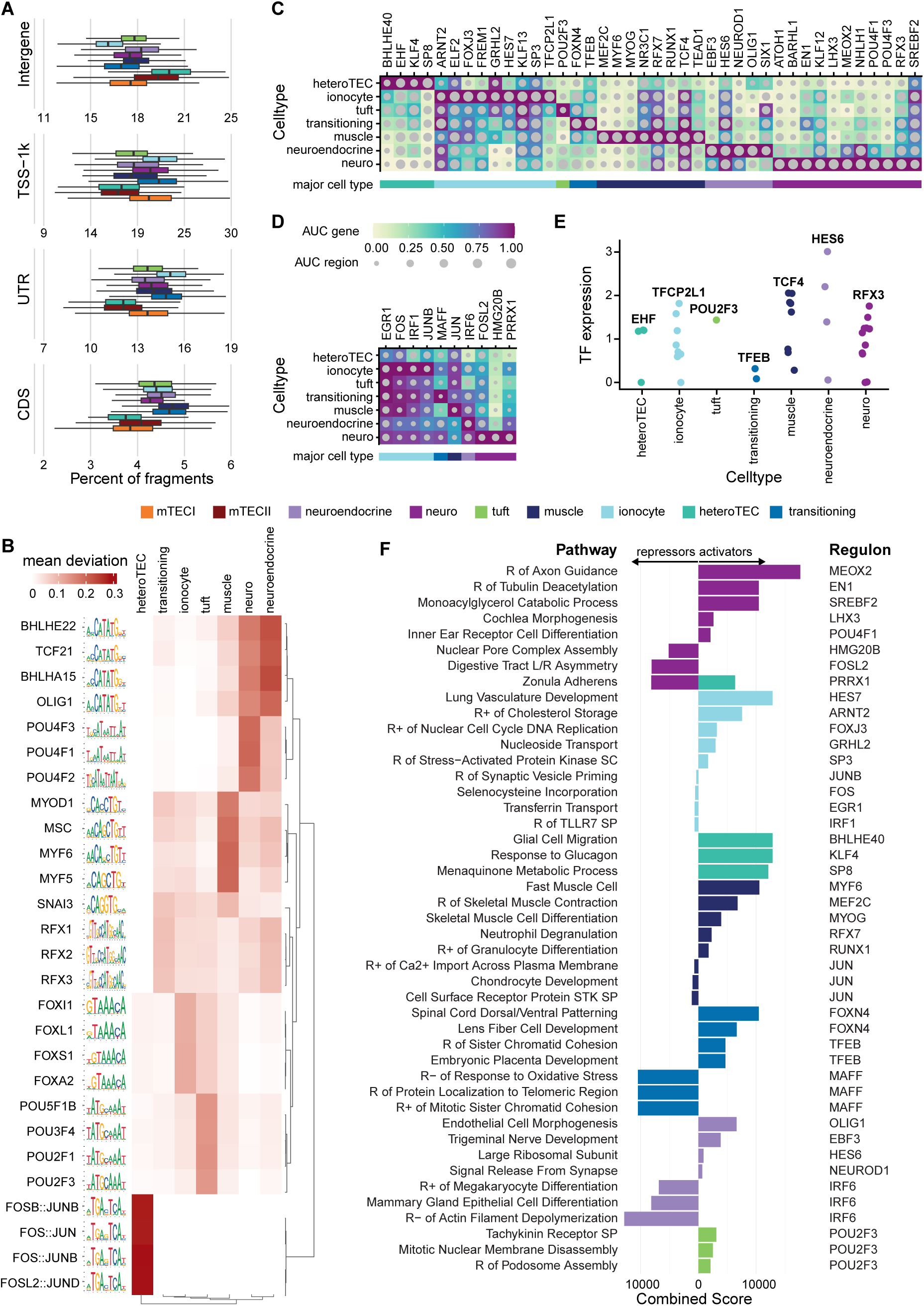
Lineage-committed TECs express peripheral cell gene signatures as part of a mixed identity. **(A)** Fragment alignment percentages from the human TEC scATAC-seq data by cell type and genomic region. **(B)** Motif enrichment via chromVAR [22] calculated as the mean deviation score of each motif for each lineage-committed TEC type; 4 motifs with the highest mean deviation score for each cell type are shown. Gene-based (color scale) and chromatinbased (size scale) Scenic+ AUC scores for the lineage-committed TEC-associated activator **(C)** and repressor **(D)** regulons. Regulons are named by their driving TFs. **(E)** Regulon-driving TF expression across TEC types. **(F)** Most enriched pathways in both the activator and repressor regulons associated with each of the lineage-committed TEC subtypes. R: Regulation; SC: Signaling Cascade; SP: Signaling Pathway.

Next, we focused on lineage-committed TECs to identify the regulatory programs underlying their commitment. Using chromVAR [22], we found that TF-motif accessibility reflected the corresponding peripheral cell programs (Figure 4B). For example, skeletal muscle TFs (*MSC*, *MYF6*, *MYF5*, *MYOD1*) [23–26] were accessible in muscle TECs, as was the tuft master regulator *POU2F3* [27] and the ionocyte TF *FOXI1* [28, 29] in tuft and ionocyte TECs, respectively.

To further link TF motif accessibility with gene expression, we jointly analyzed snRNA-seq and scATAC-seq data (Supplementary Fig 17A) for regulons that are associated with a particular TEC type [30]. We identified 95 regulons across 17 TEC types ((Supplementary Fig 17B,C), including 50 lineage-committed TEC regulons. Of these, 40 were identified as activators (Figure 4C), whereby the expression of the regulon-driving TF opens the target chromatin regions and leads to increased expression of the target genes. The remaining 10 lineage-committed TEC regulons were repressors, where expression of the driving TF leads to closing of target regions and consequently reduced expression of the target genes (Figure 4D). This integrative analysis highlighted TFs with measurable downstream consequences beyond motif presence (Figure 4B, Supplementary Fig 17D). For example, neuro TECs expressed *ATOH1* and *USH2A* (Supplementary Fig 17G), genes with established roles in brain and inner ear development [31, 32]; motifs for *POU4F3*, which mediates cochlear cilium development [33], were preferentially accessible; and the *POU4F3* regulon included both *ATOH1* and *USH2A* among its targets. Tissue-specific regulon-driving TF expression was highest in the corresponding TEC populations (Figure 4E), with *POU2F3* and *EHF* showing near-exclusive expression in tuft TECs and heteroTECs, respectively.

To interpret regulons functionally, we performed over-representation analysis on target genes associated with each regulon and corresponding lineage-committed TEC type (Figure 4F). Activator regulons were enriched for pathways matching the peripheral cell counterpart of each lineage-committed TEC: skeletal muscle contraction and differentiation in muscle TECs, axon guidance and cochlear and inner ear programs in neuro TECs; ion transport and pH regulation in ionocyte TECs [34], together with lung vasculature development and nucleoside transport, a key component of airway pH homeostasis [35]. Repressor regulons, in turn, tended to target programs characteristic of other peripheral lineages. For instance, pathways corresponding to the zonula adherens, an adhesion complex found in epithelial cells [36], were activated in heteroTECs but repressed in neuro TECs. Repressors in neuroendocrine and muscle TECs were associated with mammary gland epithelial cell development, megakaryocyte differentiation, and actin depolymerization, while muscle TEC repressors were implicated in chondrocyte development.

Taken together, lineage-committed TEC development appears to couple activation of peripheral cell-like gene expression signatures and chromatin accessibility with repression of competing lineage programs. This provides a mechanistic explanation for why TEC states can resemble specific peripheral lineages while maintaining a thymus-specific mixed identity.

### BMC-TECs exhibit progenitor-like properties

CRA module scoring of human TECs identified BMC-TECs, which expressed a stem-cell CRA module and occupied a distinct branch in the phylogeny, together suggesting a potential progenitor state (Figure 1B,C and Figure 2B). These cells are enriched for the expression of basement membrane collagens *COL4A5*, *COL4A6*, and *COL8A1*, with limited to no expression of classical cTEC (*LY75*, *PSMB11*, *PRSS16*) and mTEC markers (*KRT15*, *CLU*, *AIRE*, *HLA-DR*; Figure 5A). They express the highest level of *ITGA6* (coding for CD49f) amongst the TEC subsets, which is associated with stemness in other epithelial tissues such as skin [37], cornea [38] and gall bladder [39]. CD49f can pair with CD104 (gene: *ITGB4*) to form the *α*6*β*4 integrin, which anchors basal epithelial cells, including stem/progenitor compartments, to laminin-rich basement membranes. Both *ITGA6* and *ITGB4* were highly expressed in BMC-TECs (Figure 5A). Importantly, previous *in vitro* attempts to differentiate human induced pluripotent stem cells into TECs identified *ITGB4* as a marker expressed early in the differentiation process, implicating it as a TEC stem/progenitor marker [40]. We therefore hypothesized that these BMC-TECs were in a basement-membrane-proximal progenitor state rather than mature cTEC or mTEC identity. Moreover, BMC-TECs expressed the lowest level of CD200 relative to mature TEC subsets, indicating a CD200-population is likely to carry progenitor-like potential (Figure 5A).

**Figure 5.**
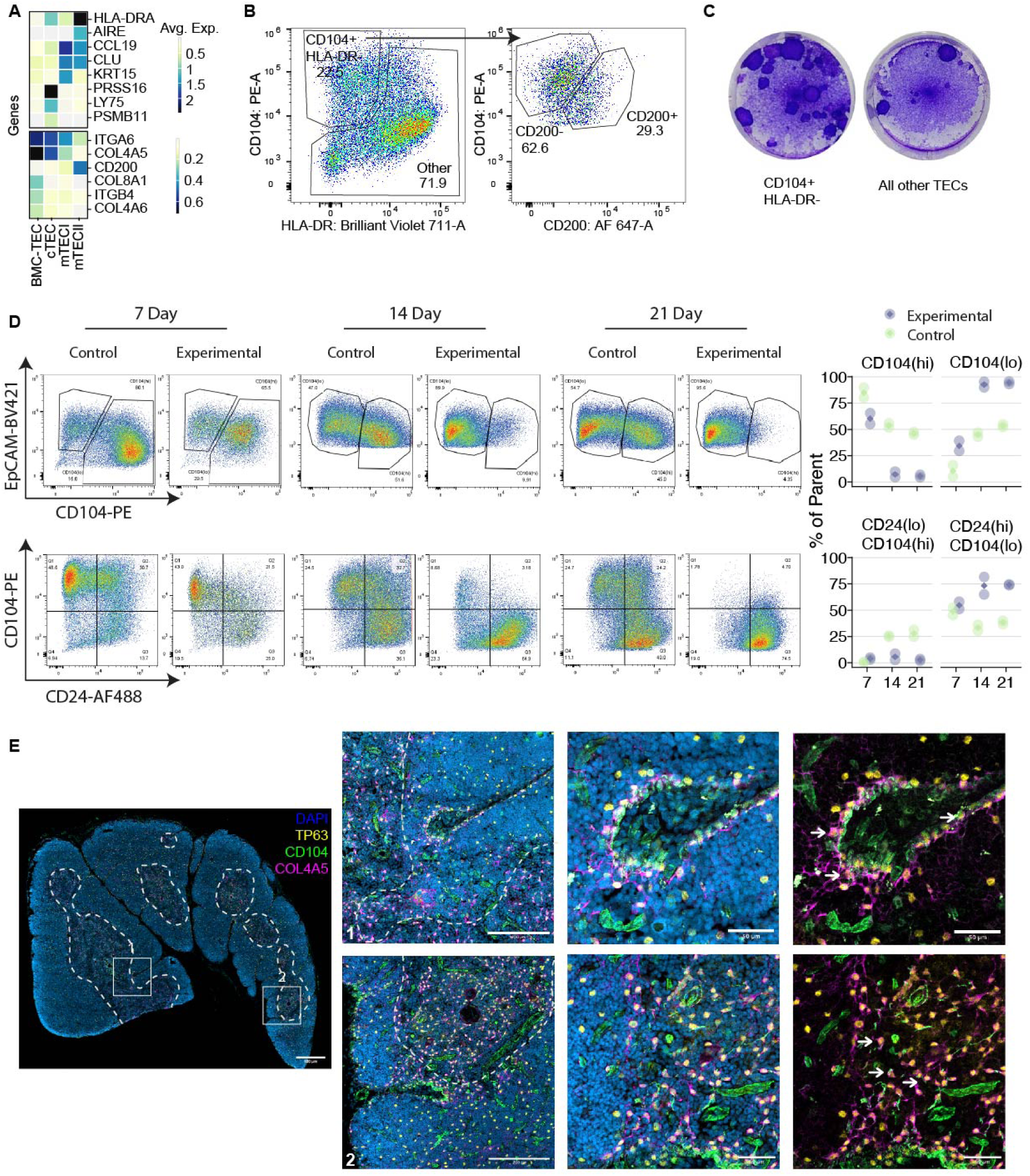
Assessment of progenitor-like traits in BMC-TECs. **(A)** Expression of select differentially expressed genes across classical TEC subsets. Top: marker genes for cTEC, mTECI and mTECII. Bottom: genes of interest relevant to the BMC-TEC subset. **(B)** Live BMC-TECs were sorted based on surface expression of CD104, HLA-DR and CD200, where BMC-TECs are HLA-DR(-/lo)CD104+CD200-. **(C)** Colony-forming potential of human TEC subsets: HLA-DR(-/lo)CD104+ and all other TECs. **(D)** Differentiation potential of sorted HLA-DR(-/lo)CD104+CD200-TECs (BMC-TECs) after three weeks of ex vivo expansion in culture on an irradiated 3T3-J2 feeder layer. Expression of EpCAM, CD104 (immature marker), and CD24 (mature marker) were used to assess the differentiation state of BMC-TECs at 7-, 14-, and 21-day time points under control or experimental (supplemented with retinoic acid and human RANKL) conditions. Representative flow plot (left) and quantification across three biological replicates (right); diamond indicates mean. **(E)** Immunofluorescence of human thymic cryosection (10 µm thick) used for spatially locating BMC-TECs at 10x magnification (scale bar: 500 µm) stained for DeltaNp63-AF647 (yellow), CD104-AF488 (green), COL4A5-AF568 (magenta); nuclei are stained with DAPI (blue). Sections imaged at 20x magnification show colocalization of DeltaNp63, COL4A5, and CD104 in perivascular spaces (1, top) and the cortical-medullary junction (2, bottom). Scale bars, 50 µm.

We translated this genomic annotation into a sorting scheme and cultured live human TECs (Figure 5B, Supplementary Fig 18A) isolated by surface expression of CD104 and HLA-DR, where BMC-TECs are CD104+HLADR-/lo, then assessed self-renewal and differentiation into mature TECs *ex vivo*. For colony-forming unit assays, we gated cells into CD104+HLADR-/lo TECs and all remaining TECs and cultured them on irradiated 3T3-J2 feeder cells. CD104+HLADR-/lo TECs formed more colonies than other TECs (Figure 5C). We then assessed differentiation using sorted CD104+HLADR-/lo CD200-TECs. Addition of the CD200-criterion removed cells potentially differentiating into more mature subsets, which could confound differentiation results. We expanded these cells on feeders for three weeks and then switched to differentiation media containing retinoic acid and human recombinant RANK-Ligand. TEC maturation was assessed weekly by flow cytometry for CD104 (higher in BMC-TECs), EpCAM (higher in mature TECs), CD24 (restricted to lineage-committed TECs), and HLA-DR (highest at the intermediate mTECII stage prior to lineage commitment, Supplementary Fig 18B) [7, 8, 40]. Over the three-week time course, we observed a continued decrease in CD104 expression with a concomitant increase in EpCAM and CD24 levels, indicating a shift from immature progenitor cells to mature differentiated cells (Figure 5D, Supplementary Fig 18C). We also observed a transient increase in HLA-DR expression in the CD24-negative cells at the 7-day time point, which disappeared at later time points. Overall, this sequential change in cell marker expression over the differentiation time course is consistent with the known mTEC differentiation trajectory: continuous increase in EpCAM and CD24 expression with transient expression of MHC class II molecules in the intermediate stage.

To investigate the tissue distribution of BMC-TECs, we performed immunofluorescence staining of human thymic cryosections using a three-marker combination: DeltaN p63 (pan-TEC marker; Supplementary Fig 19A,B), *ITGB4* and *COL4A5* (both highly expressed in BMC-TECs). We observed triple positive cells in perivascular spaces and cortical-medullary junctions, suggesting that BMC-TECs localize within these regions (Figure 5E; single-color controls in Supplementary Fig 19C,D).

In summary, genome-scale CRA module annotation, lineage tracing and integrated single-cell transcriptomic profiling nominated BMC-TECs and the surface markers needed to isolate them. Self-renewal and differentiation assays confirmed progenitor enrichment in this population, with basement-membrane-associated identity, increased colony formation, and *ex vivo* differentiation toward mature classical and subsequent lineage-committed TEC states.

### Conserved and species-specific regulation of human and mouse TECs

Mouse genetic knockout models have defined TFs that are essential for lineage-committed TEC subtypes not represented in human, including entero/hepatocyte [41] and microfold [9, 11] TECs. Recent scRNA-seq profiling of human and mouse TECs suggested conservation of lineage-committed subsets across species based on marker gene expression [8]. In addition, the abundance of subtypes that are conserved between species varied widely; tuft and muscle TECs exhibited opposite relative abundances in mice and humans [8]. Mouse lineage-committed TECs such as tuft and microfold TECs also carry out functions beyond classical tolerance induction [9, 11, 41]. Together, these observations motivated a systematic comparison of regulatory components in lineage-committed TECs from the two species based on our comprehensive census and molecular profiling of human TEC populations.

To test conservation of lineage-defining TFs and regulons, we annotated five published mouse TEC scRNA-seq datasets comprising more than 24,000 cells [5, 9, 11, 42, 43] with the same workflow used for our human TEC dataset (Supplementary Fig 20A; Supplementary Table 9). Of these, 4,725 cells from one study [11] had paired scRNA-seq and scATAC-seq profiles for regulon analysis.

The relative abundance of lineage-committed TECs varied substantially between human and mouse (Supplementary Fig 20B). We confirmed the prior observation of reversed tuft and muscle TEC abundances [8], indicating that this pattern is robust across tissue-processing protocols (Supplementary Table 5). Ionocyte TECs accounted for over 8% of human lineage-committed TECs but only 0.4% in mouse, whereas ciliated TECs were more than tenfold more frequent in mouse than in human. Our annotation of the mouse profiles did not resolve a counterpart to human BMC-TECs.

To quantify transcriptional conservation, we again used neighbor voting [20]. We found strong cross-species correspondence for classical TECs and for several lineage-committed TEC classes, notably ciliated, neuro, neuroendocrine, and muscle types (Figure 6A). Human *VIM*+ TECs, which highly expressed fibroblast and endothelial CRAs, corresponded transcriptionally to age-associated TECs previously identified in mouse [43]. Interestingly, we do not see this age association in humans, with *VIM*+ TECs found across all life stages (Supplementary Fig 20). Human heteroTECs had no direct counterpart in mice but scored highly against mouse skin, lung/goblet, entero/hepatocyte, and goblet lineage-committed TECs. We therefore re-embedded and subclustered heteroTECs (Supplementary Fig 20C) and repeated neighbor voting against the mouse skin, lung/goblet, entero/hepatocyte and goblet TECs (Figure 6B). One subcluster tracked mouse skin TECs, but the remainder still mapped to mixtures of mouse subtypes, indicating that human heteroTECs represent a mosaic state.

**Figure 6.**
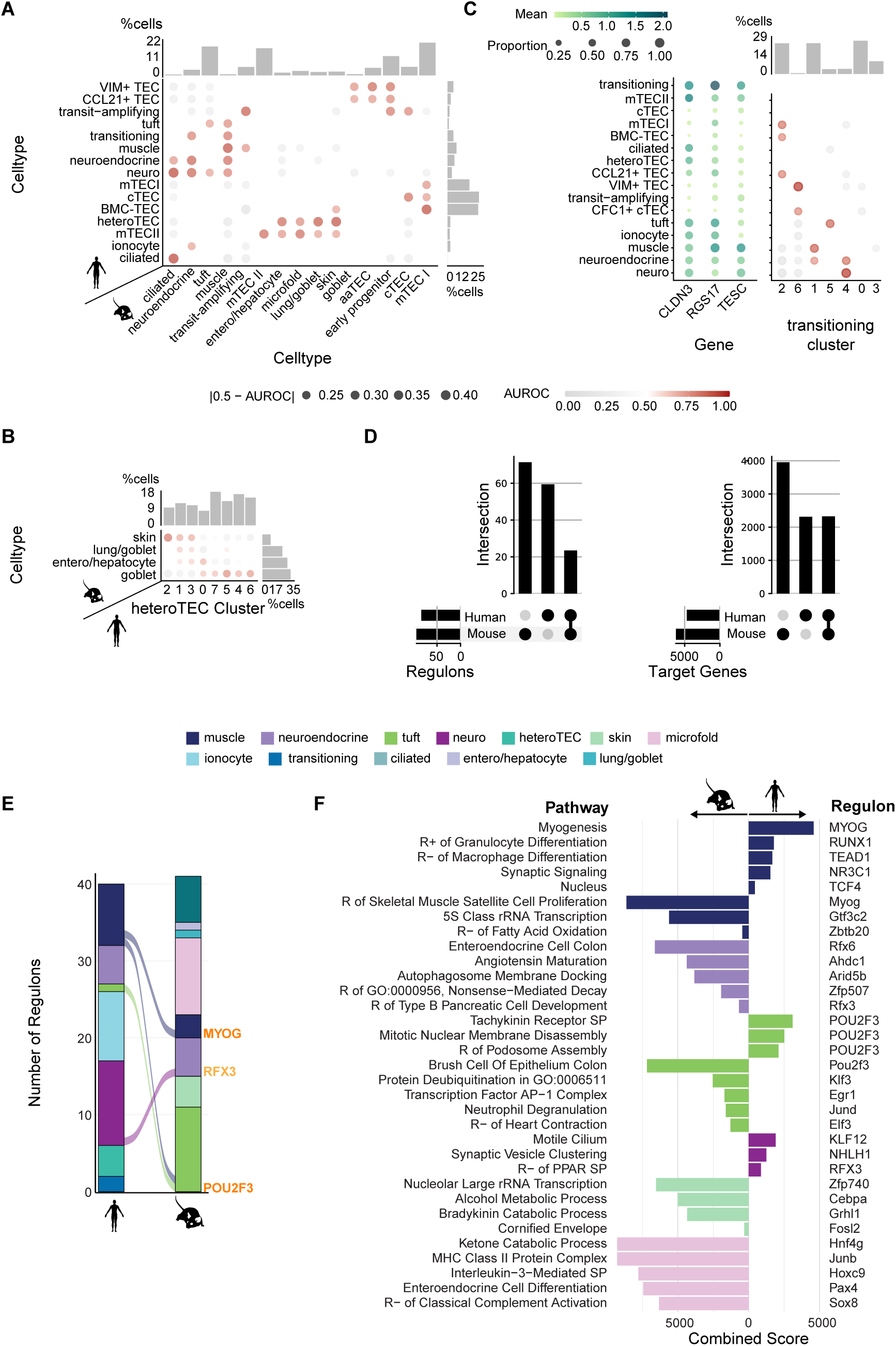
Conserved and species-specific regulation of human and mouse TECs. Panels A-C compare the integrated human TEC snRNA-seq dataset with five published mouse TEC scRNA-seq datasets; panels D-F compare regulons inferred from paired human TEC snRNA-seq and scATAC-seq data with regulons inferred from 4,725 mouse cells with paired scRNA-seq and scATAC-seq profiles. **(A)** Metaneighbor transcriptomic comparison of human and mouse cell types. **(B)** Metaneighbor transcriptomic comparison of mouse skin, goblet, entero/hepatocyte, and lung/goblet TECs and human heteroTEC subclusters. **(C)** Gene expression of human transitioning marker genes in different human TEC types (left) and their metaneighbor transcriptomic comparison to human transitioning subclusters (right). **(D)** Conserved and non-conserved (dot matrix) TF-driving regulons (left) and regulon target genes (right) in human and mouse. **(E)** Conserved regulons (ribbons and labels) and their assigned cell types in human and mouse. **(F)** Most enriched pathways in the activator regulons associated with human and mouse lineage-committed TEC subtypes.

We performed the same re-embedding, re-clustering and neighbor-voting procedure on human transitioning TECs (Supplementary Fig 20D), defined by the joint expression of the genes *CLDN3*, *RGS17*, and *TESC* (Figure 6C). Five out of the resulting seven clusters could be further resolved into individual subtypes corresponding to neuro/neuroendocrine-like, muscle-like, tuft-like, *VIM*+-TEC-like, and mTECI-like programs when compared with other human TEC types (Figure 6D). The close phylogenetic relationship between transitioning and neuroendocrine lineage-committed TECs in our somatic-variant lineage tracing (Figure 2B) is this relationship between transitioning TECs and specialized lineage-committed TECs. We did not recover a clear mouse analog in our analyses.

The neighbor-voting results showed that mouse transcriptional programs could be predictive of their human counterparts (and vice versa) despite abundance differences. To connect these similarities to mechanisms, we directly compared the regulons driving lineage-committed TEC programs in human and mouse. Analogous to the human-specific analyses (Figure 4C,D), we inferred regulons from 4,725 mouse cells with paired scRNA-seq and scATAC-seq profiles [11], yielding 95 activator regulons that were differentially active in mouse, 23 of which were driven by TFs with human orthologs that also drove human activator regulons in lineage-committed TECs (Figure 6D, left panel). Restricting to orthologous genes, we identified a total of 6,235 mouse and 4,594 human regulon targets, 2,302 of which were shared between the species (Figure 6D, right panel).

For most shared drivers, the cell types with peak regulon activity differed between species (Figure 6E, Supplementary Fig 20E, F). Exceptions included tuft (*POU2F3*/*Pou2f3*) and muscle (*MYOG*/*Myog* [44]) TECs, and *RFX3*/*Rfx3*, which linked human neuro TECs to mouse neuroendocrine TECs in a pattern consistent with known cilium-associated roles in brain development [45, 46].

Pathway over-representation analysis of conserved activator regulons reinforced peripheral cell-matched regulon signatures in each species (Figure 6F). Human muscle regulons emphasized myogenesis, whereas mouse muscle regulons highlighted skeletal muscle satellite-cell proliferation programs; mouse neuroendocrine regulons over-represented enteroendocrine and pancreatic pathways, whereas human neuro regulons emphasized cilium and synaptic vesicle genes.

Finally, we compared prerequisites for classical MHC-dependent antigen presentation. Muscle-and ciliated TEC-restricted antigen-presentation sufficed for tolerance induction in mice [9, 10], suggesting that mouse lineage-committed TECs play a role in central tolerance induction. Indeed, mouse lineage-committed TECs expressed the antigen-processing components and co-stimulatory molecules necessary for MHC-mediated antigen presentation and thymic selection (*H2-DMa*, *Cd40*, *Cd80*, and *Cd86* [47–50]), albeit at lower levels than mTECIIs (Supplementary Fig 20G, H). By contrast, across the human lineage-committed TEC subtypes, only HLA-DMA and CD40 were expressed among the mediators assessed for antigen presentation and selection (*HLA-DMA*, *CD40*, *CD80*, and *CD86* [48–51]) (Supplementary Fig 20G, H). Expression levels reached only 1–25% of *AIRE^+^* mTECII levels, with ciliated TECs (78%) and heteroTECs (83%) as exceptions. Thus, most human lineage-committed TECs lack expression of the complete co-stimulatory program observed in mouse, suggesting that their capacity for direct classical MHC-dependent tolerance induction may be limited or subtype-specific.

Overall, human and mouse TECs share transcriptomic organization and a common strategy of activating peripheral cell-type pathways. However, human mosaic states (heteroTECs, transitioning) and sparse conservation of master regulons and co-stimulatory expression point to species-specific adaptations of lineage-committed TEC identity.

## Discussion

Despite extensive single-cell profiling of the human thymic epithelium, the genomic regulatory logic by which lineage-committed TECs acquire mixed identities through the expression of peripheral cell signatures has remained unknown. Additionally, it has been unclear whether those programs are related by common ancestry and conserved regulatory components or if lineage-committed TECs arise via distinct trajectories. Our comprehensive molecular and functional analyses of human TECs yield five central insights.

First, lineage-committed TECs in humans share a common ancestor despite their divergent peripheral cell signatures. Second, these cells retain broad cross-lineage CRA expression that is largely absent from matched peripheral cell types, indicating a thymus-specific mixed identity layered on TEC differentiation. Third, paired chromatin and regulon analyses link lineage commitment to coordinated activation of peripheral cell-associated pathways together with repression of competing programs. Fourth, CRA module scores and integrated annotation identified BMC-TECs as a progenitor state, exhibiting increased self-renewal and the ability to differentiate *in vitro.* Fifth, cross-species analyses reveal shared regulatory logic but substantial divergence in key drivers, subtype abundances, and co-stimulatory programs, emphasizing limits of direct mouse-to-human extrapolation. Additionally, we have developed a new somatic-variant based method to map phylogenies in human cells using paired snRNA-seq and ATAC-seq datasets.

Lineage-committed TECs are not simple copies of their peripheral counterparts. For example, mouse thymic tuft cells express taste receptor pathways unlike intestinal tuft cells [10], and mouse microfold TECs are more similar to other TECs than to their peripheral counterparts [9]. We extend this observation to a general property of human lineage-committed TECs. Despite lineage commitment, they co-express CRA modules from multiple peripheral programs at higher levels than corresponding peripheral cells, while expressing lower fractions of corresponding CRAs. Shared ancestry suggests that peripheral programs may be progressively resolved from a common, transcriptionally broad state rather than initiated independently in unrelated cells. Muscle lineage-committed TECs have been reported to mature in stages aided by distinct transcription factors [8]; it remains to be shown whether cell-type-specific CRA expression increases while non-specific CRA expression decreases as lineage-committed TECs mature.

Our phylogenetic reconstruction supports this shared developmental origin, although it assumes that cells assigned to the same transcriptomic type share a lineage and remains sensitive to uneven coverage and missing data. Pseudo-bulking improves mutation detection but removes cell-level variation, while binary mutation calls discard information contained in allele frequencies. Future studies could improve lineage resolution by enriching individual lineage-committed populations through prospective sorting [8], using spatial ATAC-seq to distinguish progenitors across medullary niches, and developing single-cell phylogenetic models that incorporate allele frequencies.

Our systematic curation of CELLxGENE enabled us to move from the coarse-grained, tissue-level TRA framework to a cell-type-resolved CRA framework for characterizing TEC identities and states. TRAs capture ectopic expression of genes normally restricted to particular tissues, whereas CRAs resolve programs associated with specific peripheral cell types. This CRA framework revealed the mixed peripheral identities of lineage-committed TECs: lineage-committed TECs expressed fewer cell-type-restricted genes than mTECII while adopting cell-type-specific identities and retaining substantial shared CRA expression, a pattern consistent with a transition from broad antigen expression toward specialized identities. However, the CRAs were derived from a manually curated, representative downsample of CELLxGENE. The 48 categories capture broad, functionally coherent cell types but may obscure variation associated with tissue context, cell state and continuously varying subtypes [52–54]. Aligning CELLxGENE annotations with standardized cell ontologies and evaluating alternative levels of cell-type granularity should further refine the CRA framework

Whether lineage-committed TECs pass through an AIRE-expressing state and contribute directly to classical MHC-mediated tolerance remains unresolved, particularly given their limited expression of co-stimulatory molecules. Mouse lineage-committed TECs also perform functions beyond self-antigen presentation. Tuft TECs support invariant natural killer T cell development [10, 12], endocrine TECs regulate thymic involution, and microfold TECs support the generation of thymic IgA-positive plasma cells [11]. Peripheral resemblance therefore does not necessarily predict thymic function. For this reason, we use the term lineage-committed TECs rather than “mimetic cells,” which may imply that these cells simply reproduce the functions of their peripheral counterparts.

BMC-TECs provide a candidate cellular origin for human TEC diversification and a tractable population with which to test this model. Their overlap with previously described CD49f-high CD200-negative TECs [55] connects genomic annotation to an isolatable population while their colony formation and ex vivo differentiation support a progenitor role. The CD104-positive, HLA-DR-negative or low cells used in our assays were expanded on irradiated feeders in two-dimensional culture, which cannot fully reproduce the native extracellular environment. Consequently, the differentiated cells are not expected to recapitulate completely the lineage-committed states observed *in vivo*. Likewise, microscopy identified BMC-TECs using DeltaNp63, ITGB4 and COL4A5, but ITGB4 and COL4A5 are expressed at lower levels in several other TEC populations. Their colocalization therefore enriches for, rather than uniquely identifies, BMC-TECs. Determining whether these cells generate the full range of TEC states *in vivo* and how their basement-membrane niche influences fate will require more specific markers and lineage-resolved studies in human tissue.

Cross-species comparison distinguished conserved regulatory principles from species-specific states. Divergence in most candidate drivers, cell-state abundance and co-stimulatory expression suggests that similar peripheral programs may support different thymic functions in mice and humans. Mouse models therefore remain informative for conserved mechanisms, but human tissue will be required to determine the functions of human-specific and mixed TEC states. Regulon-inference methods also recover activators more effectively than repressors [30], and we identified no repressor regulons in mouse lineage-committed TECs. This absence should not be interpreted as evidence that repression does not occur in these cells. Larger paired transcriptomic and chromatin-accessibility datasets will be needed to compare repressor programs across species with greater power.

Together, these findings establish lineage-committed TECs as thymic epithelial states whose mixed peripheral programs arise through shared ancestry and coordinated gene regulation rather than as simple copies of peripheral cells. This framework shifts the central question from which peripheral identity each TEC resembles to how developmental history and regulatory circuitry shape its thymic function. Because TEC composition changes with age in humans and mice [6, 42, 43], larger post-pediatric cohorts should determine how ageing affects CRA expression and regulation. Ultimately, defining which human TEC states present antigen and which support other thymic functions will be necessary to understand how epithelial diversity sustains immune tolerance and whether its disruption contributes to autoimmunity.

## Methods

### Cell type specific gene expression

#### CELLxGENE data processing

The CELLxGENE database includes single-cell RNA-seq data from over 479 different single-cell RNA sequencing datasets with 61,888 genes and 62,634,126 cells (census version 2025-01-30), with harmonized and standardized metadata for each dataset. We initially downloaded the metadata for all cells in CELLxGENE census and applied a cell and tissue type harmonization, quality control and downsampling for downstream analyses.

#### Harmonization of Cell and Tissue Types

CELLxGENE metadata contains annotations for the study tissue (column: *tissue*, *tissue_general*) and author-annotated cell types (column:*cell_type*). We curated these tissue and cell types to enable harmonized analyses focusing on shared cell types across tissues. Supplementary Table 2 and Supplementary Table 1 show the raw and curated cell and tissue types. Imprecise tissue types (e.g. ‘Neck‘) and cell types (e.g. ‘cell‘, ‘blood cell‘) were removed for downstream analyses and are marked as ‘QC_fail’. For all downstream analyses, we consider the pair of a specific cell type in a specific tissue a category, i.e., basal_epithelium-breast, myoepithelium-breast, and myoepithelium-salivary_gland are three distinct categories.

#### Data download and curation

We filtered cells based on the following criteria in the metadata sheet: i) No duplicate cells were allowed, ii) assays were 10x 3’ transcription profiling, 10x 3’ v1, 10x 3’ v2, 10x 3’ v3, 10x 5’ transcription profiling, 10x 5’ v1, and 10x 5’ v2, iii) non-diseased donors, iv) only primary cells, v) no donors with unknown development stage and vi) expression of at least 200 genes. In addition, we removed cells with ‘QC_fail‘ in the harmonization steps (Supplementary Fig 1A). Down-sampling analyses on a subset of categories types showed a minimum of 50 cells per category were required for reproducible results on pseudobulk level (with a mean *r*^2^ = 0.9685, Supplementary Fig 1B), thus any category with fewer than 50 cells was removed. With 5000 cells per category, we obtained a mean *r*^2^ = 0.9995, choosing this as the upper bound needed for consistency, while making computation feasible. To account for additional downstream filtering based on QC criteria not present in the metadata, i.e., percentage of mitochondrial gene expression, we sampled a random subset of 8000 cells (or less if the category included fewer cells) indexed by their soma_joinid from each category and used the API to download the corresponding gene expression data. After additional filtering on mitochondrial percentage less than 15%, all categories were downsampled to an upper bound of 5000 cells. The final dataset contained 1,558,164 cells with 61,888 genes for 580 categories with 45 tissues and 48 distinct cell types.

#### Cell-type-restricted antigens

For each of the 48 distinct cell types in the curated and QC dataset we computed genes that are specific to this cell type, using the *τ* metric [56] previously applied [3] and benchmarked [57] for obtaining tissue-specific antigens (TRAs). We thus named the set of genes specific to each cell type cell-type-restricted antigens (CRAs). As input to calculating *τ*, we calculated the pseudo-bulk expression for each cell type across tissues by summing counts for each gene across all cells and dividing the final gene expression by the total number of counts per cell type. We then multiplied this normalized count by 10,000 to obtain gene expression as counts per 10,000 (CP10K). Genes which had a maximal expression of < 1*CP* 10*K* were removed prior to calculating *τ*. *τ* was calculated as

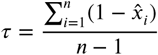

where

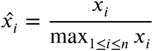

and *x_i_* is the gene expression in cell type *i*. Based on the saddle point at *τ* = 0.73 (Supplementary Fig 1C), we considered genes with *τ* ≥ 0.73 as CRAs. We then assigned the cell type specificity using the binarization approach described in [56]: for each gene with *τ* ≥ 0.73, we sort the gene expression values across cell types and compute the distance between neighboring gene expression values in this sorted vector. We find the maximal distance in this vector and choose the expression value at the right hand side of the maximal distance as the threshold above which cell types are considered to specifically express the gene (Supplementary Fig 1D).

### Tissue processing

#### Tissue samples

Discarded thymus tissues obtained from human donors undergoing medically necessary surgery were collected under the Mount Sinai Biorepository with a Waiver of Consent. Additional tissue was collected under the Northwell Health Biospecimen Repository (IRB 20-0150) and New York University Langone Health (i25-00159_MOD07) with patient consent. The use of these human tissues was reviewed by the Cold Spring Harbor Institutional Review Board.

#### Tissue digestion

Human thymus samples were delivered to the lab for processing within 3 hours post-surgery. Up to 2g of the tissue was taken for isolation of TECs (Supplementary Fig 2A). Tissue was placed into a dish containing tissue buffer (RPMI1640, 5% FBS, 1x Pen/Strep, 10mM HEPES) and cut into progressively smaller pieces. Thymocytes were released from tissue by stirring tissue pieces for 10 in 10ml of cold tissue buffer per 1g of tissue on magnetic stirrer. The remaining tissue was placed in RPMI1640 media containing a mixture of digestive enzymes (Collagenase IV 0.6 mg/ml, Dispase I 1.6 units/ml, and Dnase I 80 units/ml) and digested for 25 at 37°C to release TECs. This digestion step was performed for up to three rounds. A final round of digestion using TrypLE for 5 at 37°C was done to break up microscopic clumps of cells into single cells. The released cells were washed with tissue buffer after each round to deactivate the enzymes.

#### Epithelial cell enrichment by panning

Polystyrene petri dishes were coated with goat anti-mouse IgG antibodies in Tris pH 9.5 at a concentration of 100µg of antibody per 4ml of Tris pH 9.5 per petri dish for at least an hour at room temperature. After coating, dishes were washed three times with 1xPBS and kept in 1xPBS until ready to use. Post-digest cells were stained with a cocktail of unconjugated antibodies against immune cells and red blood cells (mouse anti-human CD3, clone UCHT1; mouse anti-human CD4, clone RPA-T4; mouse anti-human CD8a, clone RPA-T8; mouse anti-human CD45, clone HI30; mouse anti-human CD235a, clone HI264; all purchased from BioLegend) for 30 on ice, washed and resuspended in cell enrichment buffer (1xPBS, 4% FBS, DNase I). Cell suspension was dispensed (‘panned‘) onto the anti-mouse IgG coated petri dishes at a concentration of 150 million cells per petri dish. Supernatant containing TECs was collected after 20 of incubation at room temperature. The panning process was repeated for a total of three rounds. An optional fourth round of panning was performed if the first three rounds were deemed insufficient.

#### Live sorting TECs

After enrichment by panning, the remaining cells were stained with EpCAM-BV421 antibody (clones 9C4 and CO17-1A, BioLegend), HLA-DR-BV711 antibody (clone L243, BioLegend), CD104-PE antibody (clone 58XB4, BioLegend), CD200-AF647 (clone OX-104, BioLegend), and a viability dye (Fixable Near-IR, Invitrogen) at a concentration of 1 million cells per 100µl for 30 minutes on ice. After staining, cells were washed twice with enrichment buffer at 350g for 5 minutes at 4°C and resuspended in enrichment buffer. Cells were sorted using the Sony SH800 Cell Sorter with an 100um nozzle in Targeted Mode into TEC media supplemented with an additional 10% of heat-inactivated FBS (Supplementary Fig 2A). The gating strategy was as follows: first gate on all cells except debris, then gate on EpCAM+ non-autofluorescent cells, remove dead cells, gate on singlets, and draw gates for CD104+ HLADR-/lo TECs and all other TECs; within CD104+ HLADR-/lo TECs, gates were drawn for CD200-and CD200+ subsets. On the SH800 sorter, the unused FITC detector was activated for detecting autofluorescence signals given off by the cells. For colony-forming unit assays, CD104+ HLADR-/lo TECs and all other TECs were collected as two populations. For differentiation assays, CD104+ HLADR-/lo CD200-TECs were sorted based on enrichment of BMC-TECs within this fraction in the integrated scRNA-seq dataset.

#### Colony-forming unit assay

For the colony-forming unit assay, 1 cryovial of irradiated 3T3-J2 fibroblast feeder cells (STEMCELL Technologies) was thawed and split equally into 2 wells of a 6-well tissue culture plate. The feeder cells were cultured in DMEM/F-12 medium supplemented with 10% heat-inactivated FBS, 1x Glutamax, 1x Pen/Strep for at least 8 hours prior to sorting of the TEC subsets described above so the layer can attach to the well. 1,000 cells from each of the two sorted populations (CD104+ HLADR-/lo TECs and all other TECs) were seeded onto the feeder layer in their respective wells and cultured in TEC media for up to 14 days at 37°C. Half of the tissue culture media was changed 3-days after the initial seeding of the TEC cells, then every other day. At the end of the culture duration, cells were washed once with 1xPBS then fixed with 4% paraformaldehyde solution for 25 minutes at room temperature. After fixation, the paraformaldehyde solution was removed and cells washed once with 1xPBS to remove residual paraformaldehyde. Cells were then stained with a 0.05% w/v solution of Crystal Violet at room temperature for 15 minutes, washed with 1xPBS, and cell colonies imaged.

#### Differentiation assay

Based on integrated scRNA-seq annotation, which indicated that BMC-TECs were enriched within the CD200-fraction of CD104+ HLADR-/lo cells, differentiation assays used sorted CD104+ HLADR-/lo CD200-TECs. 3T3-J2 irradiated feeder cells (STEMCELL Technologies Cat. 100-0353) were thawed in a waterbath at 37°C. Cells were suspended in 8mL DF10 media, which contains DMEM/F-12 (Gibco Cat. 21041025), 10% Fetal Bovine Serum (Corning Cat. 35-010-CV), 1x Pen/Strep, 1x GlutaMAX (Gibco Cat. 35-050-061), per 2-3 x 10 3T3-J2 cells. Cells were washed once by spinning down at 4°C, 300g, for 10 minutes. Cells were resuspended in DF10 for 1-1.5 10 cells/4mL. 4mL of the cell suspension was plated per flask, and cells adhered for a minimum of 8 hours in an incubator at 5% CO2 and 37°C before plating of TECs. Sorted CD104+ HLADR-/lo CD200-cells were resuspended at 2,500 cells/ 4mL of expansion media containing DMEM/F-12 (Gibco Cat. 21041025), DMEM (Gibco Cat. 31053028), 10% Heat-inactivated Fetal Bovine Serum (Corning Cat. 35-010-CV), 1x Pen/Strep, 1x GlutaMAX (Gibco Cat. 35-050-061), 1x HEPES (Gibco Cat. 15-630-080), 5 µg/ml Insulin (Gibco Cat. 1258501), 0.4 µg/mL Hydrocortisone (STEMCELL Technologies Cat. 7925), 0.1nM Cholera Toxin (Sigma-Aldrich Cat. C8052), 10ng/ml Human EGF, Animal-Free Recombinant Protein (PeproTech Cat. AF-100-15-100UG). DF10 media was removed from flasks and replaced with 4mL of cell suspension in expansion media. Cells were grown in expansion media for 19 days, before being transferred to differentiation media, containing; DMEM/F-12 (Gibco Cat. 21041025), 1x GlutaMAX (Gibco Cat. 35-050-061), 1x HEPES (Gibco Cat. 15-630-080), 1x Primocin (InvivoGen Cat. ant-pm-2), 1x B27 supplement with vitamin A (Gibco Cat. 17504044), 0.2 µg/mL Human TRANCE (RANKL) Animal-Free Recombinant Protein (PeproTech Cat. AF31001100UG), and 0.001mM Retinoic Acid (Sigma Aldrich Cat. R2625-500MG). Half the volume (2mL) of expansion media was changed every other day. Control groups were not transferred to differentiation media, but stayed in expansion media for the entirety of the experiment. Cells were collected 7, 14, and 21 days after adding differentiation media.

#### Cell collection

At the appropriate time point, flasks were removed from the incubator one at a time for cell collection. Media was removed, and cells were washed with 10mL of PBS pH 7.4 (Gibco Cat. 10-010-049) before trypsination with 400µL of TrypLE Express (Gibco Cat. 12-604-013) and returned to the incubator. Every 30 seconds the flask was removed, agitated, and placed back in the incubator, for a total of 5 minutes. Cells were then resuspended in 4mL of FACS buffer consisting of PBS pH 7.4 (Gibco Cat. 10-010-049), 2% Heat-inactivated Fetal Bovine Serum (Corning Cat. 35-010-CV), 2mM EDTA, 0.05%w/v sodium azide and immediately spun down for 5 minutes at 350g, followed by resuspension in 5mL of FACS buffer. This was repeated 4x per flask. After all cells were collected, cells of the same flask were spun down at 350g and 4°C for 5 minutes, and pooled together. 1 million cells were stained for flow cytometry.

#### Flow cytometry

Samples were washed once by spinning down at 350g for 5 minutes at 4°C, and resuspended in 3mL of HBSS, no Ca, no Mg, no phenol red (Gibco Cat. 14175103). Samples were then spun down at 350g for 5 minutes, and resuspended in 250µL of Fc block (eBioscience™CD16/CD32 Monoclonal Antibody Clone 93) at 1:10 concentration with HBSS (Gibco Cat. 14175103), and left to incubate at 4°C for 10 minutes. After blocking, 250µL of the staining master mix was added to the samples, and incubated for 30 minutes at 4°C. Samples were prepared for flow cytometry and stained using: LIVE/DEAD™ Fixable Blue Dead Cell Stain Kit at 1/1500 dilution (Invitrogen Cat. L23105), Brilliant Violet 421™ anti-human CD326 (EpCAM) Antibody Clone COI7-IA at 1/200 dilution, Brilliant Violet 421 anti-human CD326 (EpCAM) Antibody Clone 9C4 at 1/200 dilution, Brilliant Violet 711™ anti-human *HLA-DR* Antibody clone L243 at 1/800 dilution, Alexa Fluor 647 anti-human CD205 (DEC-205) Antibody Clone HD30 at a dilution of 1/200, Alexa Fluor 647 CD205 clone HD83 at a 1/200 dilution, Alexa Fluor 488 anti-human CD24 Antibody Clone ML5 at a 1/200 dilution, and PE anti-human CD104 Antibody Clone 58XB4 at a 1/300 dilution (all antibodies purchased from Biolegend, unless otherwise specified; CD205 HD83 was conjugated using the Invitrogen Antibody Labelling Kit Cat. A20186). After staining, the volume was brought up to 3mL with FACS buffer and spun down at 350g for 5min at 4°C. The samples were washed once with 3mL HBSS and spun down at 350g for 5 minutes at 4°C. The samples were fixed using 2% PFA (Electron Microscopy Sciences Cat. 15710) for 30 minutes at 4°C. After fixing, samples were washed 3x with HBSS (Gibco Cat. 14175103), and spun down at 550g for 5 minutes at 4°C. Samples were stored in FACS buffer at 4°C until ready for flow cytometry. Samples were run using the BD LSRFortessa Cell Analyzer within a week of preparation. Samples were filtered through a 100um filter right before analyzing on the flow cytometer. Data was analysed using FlowJo v10.10.1 software. The initial gating strategy starts with FSC-AxSSC-A to gate on the target cell population, followed by, FSC-AxFSC-H, deduced to identify and exclude doublet-cells from the analysis. To refine the final population of cells used for this analysis we incorporated an additional singlet gate, SSC-AxSSC-H. Next, we gated on the live population of cells using the ViabilityxEpCAM plot. Lastly, we included a BV605(open channel)xEpCAM gate to exclude autofluorescent cells. The population derived from these ‘quality control’ plots, is the population used for both qualitative and quantitative analysis.

#### Tissue localization of BMC-TECs by Immunofluorescence

10 µm-thick fresh frozen thymic tissue sections (stored at -80°C) were fixed with 4% Paraformaldehyde (Electron Microscopy Sciences cat. 15710-S) for 5 minutes at room temperature, followed by 3 washes with 1xPBS at room temperature for 5 minutes each. Slides were then fixed for 10 minutes at -20°C using pre-chilled acetone. Slides were removed and dried in open air for 5-10 minutes. After fixation, slices were washed again using 1xPBS with 0.02% Tween-20 (PBST) 3 times for 5 minutes each at room temperature. The tissue was then blocked for 30 minutes in a humidity chamber at room temperature with 2.5% donkey serum (Jackson ImmunoResearch cat. 017-000-121) in PBST. Primary antibody staining was performed overnight at 4°C in a humidified chamber with rabbit anti-human DeltaN p63 (1:200, Cell Signaling Technologies cat. 67825T), followed by washing with PBST, and subsequent secondary antibody staining was performed using the Alexa Fluor 647 AffiniPure Donkey Anti-Rabbit antibody (JacksonImmunoResearch cat. 711-605-152) for 1 hour at room temperature in a dark humidified chamber. Slides were washed using PBST 3 times for 5 minutes each at room temperature. The COL4A5 antibody (1:200, Thermofisher cat. PA5-49913) was conjugated using the Zenon AF568 Conjugation kit (Thermofisher cat. Z25306) according to the manufacturer’s protocol. Once conjugated, mouse anti-human CD104 Alexa Fluor 488 (1:200, R&D Systems cat. FAB4060G) and rabbit anti-human COL4A5 antibodies were used to stain the tissue in blocking buffer containing 2.5% mouse serum (Jackson ImmunoResearch cat. 015-000-120) and rabbit serum (Jackson ImmunoResearch cat. 011-000-120) in PBST for 1 hour at room temperature in the humidified chamber. A final washing step using PBST was performed 3 times for 5 minutes each at room temperature. DAPI was prepared at a 1:2500 dilution in 1xPBS, mixed overnight, and then filtered through a 0.22 µm filter. DAPI staining was done in a dark, humidified chamber for 15 minutes at room temperature. After staining, residual DAPI was washed away with two 5-minute washes of 1xPBS at room temperature. Slides were mounted with Prolong Gold Antifade (Thermofisher cat. P10144) and cured for 24 hours before imaging. The images were acquired on the Zeiss Airyscan LSM 910 microscope and analyzed using Fiji version 2.16.0/1.54p.

### Sequencing

#### 10x multiome (scRNA-seq and scATAC-seq) preparation

Enriched TECs were stained with antibodies against human EpCAM (clone 9C4 and CO17-1A, from BioLegend) and CD45 (HI30, from BioLegend) and a SYTOX viability dye (from Invitrogen). Cells were then resuspended in 500µl of nuclease-free 1xPBS containing 2% heat-inactivated FBS. Live EpCAM+CD45-cells (Supplementary Fig 2B, C) were sorted into nuclease-free 1xPBS containing 10% heated-inactivated FBS in a BSA fraction V coated 15ml falcon tube on a Sony SH800 sorter using a 100µm nozzle in Targeted mode. Sorted cells were kept cold on ice and submitted to the single-cell sequencing core facility immediately for the 10x multiome library preparation and sequencing. Of note, the compensation values were not applied to the data during the sorting of TECs from sample HT8, making the EpCAM+ cells appear positive for the negative markers in the FACS flow panel (Supplementary Fig 2B). A mixture of non-TEC cells was sorted out along with the TECs as a result, but we were able to distinguish and filter out these contaminating cells based on their gene signature in the scRNA-seq data. The compensation values were applied to all subsequent samples (Supplementary Fig 2C).

#### Parse WTv2 scRNA-seq preparation

A modified version of the standard Parse protocol (Parse fixation protocol, October 2022 version) was used for preparing cells for sequencing using the Parse Biosciences technology. Enriched TECs were stained with antibodies against human EpCAM (clone 9C4 and CO17-1A, from BioLegend), CD45 (clone HI30, from BioLegend), and a fixable Near-IR viability dye from Invitrogen (EpCAM+CD45-; Supplementary Fig 2D) for 30 on ice. After staining, the cells were washed once with nuclease-free 1xPBS at 300g for 5 at 4°C to remove as much EDTA, DNase, and RNase as possible and resuspended in 800µl of cell prefixation buffer containing 0.5% w/v BSA fraction V. The resuspended cells were filtered through a 70µm nylon filter into a BSA fraction V coated 15ml falcon tube. 250µl of cell fixation solution was immediately added to the cell suspension and mixed gently by pipetting up and down exactly three times with a p1000 pipet set to 250µl and placed on ice to fix for 10. After fixation, cells were washed twice with nuclease-free 1xPBS at 400g for 5 at 4°C to remove as much fixative as possible and then resuspended in nuclease-free 1xPBS at 1 million cells per ml for sorting. After sorting, cells were spun down and supernatant removed until about 1ml remained. To that 1ml of residual supernatant, 80µl of cell perm solution was added, mixed, and cells incubated for 3 on ice. 4ml of cell neutralization buffer was added to the cells and gently mixed with a p1000 pipet. Cells were spun down, resuspended in 150µl of cold cell buffer, and filtered through a 70µm nylon strainer into a 2ml nuclease-free tube. 2.5µl of DMSO was added to the cell suspension and mixed by gently flicking the tube three times. The DMSO addition was repeated two more times for a total of 7.5µl of DMSO and the final cell suspension was frozen down in a -80°C freezer using a Mr. Frosty slow freezing unit. The standard Parse WTv2 protocol was followed for library construction, and submitted for sequencing.

### Single-cell RNA-seq analysis of TECs

Publicly available thymic single-cell datasets provided a useful resource toward this study’s goal of characterizing lineage-committed TECs in humans and mice. However, these and our datasets have been processed using 5 different single-cell protocols: 10x 3’ V2 and V3, 10x 5’ V2, 10x multiome V1, Parse Bioscience WTv2, and SMART-seq2. Moreover, the count matrices generated in the original studies use several different versions of (often proprietary) tools. These different tool versions utilize different algorithms for analysis, which introduces an unnecessary source of variation for our pooled analysis of all datasets [58]. Therefore, we re-processed all datasets from the original FASTQ files through a common pipeline to make them as uniform as possible.

#### Data download

The FASTQ files for all datasets were downloaded from public repositories indicated in Supplementary Table 5. Metadata for each dataset (study, sample, individual, species, subspecies, sex, age, sort type, rna seq protocol) in Supplementary Table 6.

#### Reference genome and annotation

Human samples were aligned to GRCh38 (primary assembly) using Gencode annotation version V44. Mouse samples were aligned to GRCm39 (primary assembly) using Gencode annotation version vM33.

#### Read alignment, cell filtering, and count matrix generation

We processed all datasets using STARsolo (2.7.11a) [59]. STARsolo is designed for all steps of scRNA-seq data processing that include read alignment to the reference genome, assigning aligned reads to individual genes to generate the count matrix, and cell filtering to retain single cells (which depending on the sequencing protocol could be removing empty droplets and doublets, or removing cells with too few gene counts). The parameters used for STARsolo (both reference genome indexing and read alignment) for each dataset are listed in Supplementary Table 7. Different library preparation methods necessitate the use of STARsolo in slightly different manner. We used STARsolo as follows:

- **10x 3’ and multiome datasets:** Reads were aligned using the standard parameters for 10x 3’ datasets (listed in supplementary table 2). We used the EmptyDrops_CR option to call singlet cells and the GeneFull_Ex50pAS gene model using the expectation maximization (EM) option to resolve multi-mapping reads. STARsolo generates only the “raw” version of the EM count matrix that contains the gene counts for all possible 10x barcodes. The “filtered” count matrix containing only singlet cells was generated by using the filtered barcodes to subset the raw EM count matrix, thereby retaining just the required cells.
- **10x 5’ datasets:** The 10x 5’ datasets were aligned identical to the 10x 3’ dataset with the only difference being these the reads are mapped to the reverse strand of the genome.
- **Parse datasets:** Parse WTv2 is a whole-transcriptome protocol that uses a poly-T primer to capture the 3’ end of a transcript and a random hexamer primer to capture the internal regions of a transcript. We used reads originating from just the poly-T primers to keep the analysis as consistent as possibles with the 10x data. This processing procedure is described as a part of the documentation for its_a_scam (https://its-a-scam.readthedocs.io/). We selected all cells that had a read count of at least 500 for downstream analysis.
- **SMART-seq2 datasets:** The reads were aligned using the standard parameters for the SMART-seq2 dataset.

We used the default cell filtering option in STARsolo for the SMART-seq2 dataset and the GeneFull_Ex50pAS gene model was used without the EM option (since the SMART-seq2 protocol does not use UMIs).

### Quality control of scRNA-seq data and selection of TECs

#### Quality control for scRNA-seq data

Single-cell RNA-seq and single-nuclear RNA-seq data were processed via scanpy [60] (version 1.9.3). Cells with a mitochondrial gene percentage greater than 15% and with fewer than 200 expressed genes were removed from downstream analysis.

#### Normalization

We filtered zero-count genes by dataset followed by filtering all datasets to the remaining genes common between the datasets and removed mitochondrial and ribosomal genes. We then normalized the datasets individually via sc.pp.normalizetotal with a targetsum = 1e4. We performed log normalization with sc.pp.log1p, combined the normalized datasets and selected the the top 3,000 highly-variable genes via sc.pp.highlyvariablegenes.

#### Initial dataset integration

We integrated the datasets using the normalized counts via scVI [61, 62] (version 1.0.3) with three categorical batch covariates: sequencing protocol, donor, and study. The scVI integration generated an integrated embedding of all the datasets that was used for downstream analysis (Supplementary Fig 6B). The Yayon dataset (Supplementary Fig 6A) integrated poorly with the other datasets and was thus excluded from further analysis. We evaluated the success of our integration procedure using the local inverse of Simpson’s Index (LISI) metric [63]. The LISI metric estimates the number of cells that can be sampled at random in a local neighborhood before selecting two cells from the same batch. Thus, this “integration LISI” (iLISI) score estimates the number of batches in the given local neighborhood. A successful batch integration aims to harmoniously merge the datasets and will therefore result in an increase in the iLISI score post-integration. We also calculated a LISI score applied to cell types instead of batches, called the “cell type LISI” (cLISI). A successful integration will yield a merged dataset where cells of the same type cluster together within local neighborhoods, so the cLISI score will approach one post-integration. We conducted this analysis prior to cell type annotation. We therefore identified mock marker-based cell type populations to be used in the cLISI calculations using previously-identified marker genes (Supplementary Table 8. Muscle cells were defined as cells having normalized expression of *MYH3* > 2.5, neuro cells with *USH2A* > 2.5, tuft cells with *POU2F3* > 2.5, cTEC’s with *TBATA* > 2.5, heteroTECs with *SPINK5* > 3, and mTECIIs with *AIRE* > 2.5 (Supplementary Fig 6D). All LISI scores were computed using the lisi package [63] (version 1.0) (https://github.com/immunogenomics/LISI).

#### TEC identification

We first used the integrated scVI embedding to identify TECs as all datasets contained non-TECs. The exception to this protocol was the Park dataset [5], as due to its size and large number of non-TECs, we only used the cells previously-identified as TECs for this analysis. We defined cell clusters using sc.pp.neighbors and sc.tl.leiden on the integrated embeddings generated via scVI and identified two distinct subsets of non-epithelial cell populations, endothelial cells (expression of *VWF* (Von-Willibrand Factor)) and T cells (expression of *CD8*, *CD4*, and *CD3*). We removed both non-TEC cell populations from downstream analysis. We identified thymic epithelial cells based on the positive expression of *EPCAM*, *KRT5*, *KRT8*, and *KRT15*.

#### TEC re-normalization, re-integration, and annotation

We extracted all cells identified as TECs into a final dataset, which we then re-normalized and re-integrated with scVI according to the procedure outlined above. We annotated the integrated TECs using an iterative clustering procedure. We defined clusters using sc.pp.neighbors and sc.tl.leiden on the integrated embeddings generated via scVI, and then annotated the clusters based on their expression of both known TEC marker genes and the CRA signatures we identified above.

We first computed CRA modules scores from the normalized counts via scanpy.tl.score_genes. For each cell, we kept scores greater than zero and normalized score values by the maximal observed score; the resulting score densities are depicted in Supplementary Fig 1E. For each cell, we classified each module as expressed if the observe score was greater than 50% of the maximal score for this module. We obtained the mean score for each module and cluster and calculated MetaNeighbor as the fraction of cells in the cluster considered to express the module over the total number of cells in the cluster. For each cluster we chose the top scoring 30% of modules (7 modules) and for each module the top 20% scoring clusters (6 clusters) for annotation. For visualization, we used hierarchical, complete linkage clustering on Euclidean distances for both rows (CRA modules) and columns (cell types).

After the initial round of TEC annotation was complete, we integrated the data in a semi-supervised manner via the scANVI model [64], which updated the scVI embeddings based on the input cell types. We then completed further iterations of clustering and annotation using the scANVI embeddings.

Some clusters exhibited elevated expression of either a single set of TEC marker genes or of a single CRA module; we annotated such clusters as consisting solely of this corresponding cell type. Other clusters expressed elevated levels of multiple sets of marker genes or CRA modules; we further sub-clustered these cells through several iterations until we resolved them into smaller clusters that exhibited elevated expression of only a single set of TEC marker genes or a single CRA module.

Jupyter notebooks containing the code for this annotation process are available at https://github.com/meyer-lab-cshl/ intecgrated_2026.

### Gene lists

We defined human protein-coding (PC) genes as the 20,070 genes that are classified as protein_coding in the gene_type tag of the Genocde GTF file (V44). We used the list of human house-keeping (HK) genes as defined by Eisenberg & Levanon; we mapped the gene identifiers from refseq to ensemble gene identifiers [4], yielding 4,211 unique genes [65].

### Robustness of expressed gene counts to cluster annotations

We determined the number of expressed genes for each cell type by pseudo-bulking the cells within each cluster (analogous to a differential gene expression analysis using single-cell data). However, since cell types are defined by first clustering the cells, and gene counts are determined from clustered cells, misclassifications of a “true” cell identity (i.e assigning a cell to the wrong cluster) will lead to an over-estimate of the number of genes expressed. For example, if a true neuro TEC was misclassified as a muscle TEC, it would lead to an over-estimate of number of genes expressed in muscle TECs. We determined the impact of any mis-clustering of cell types on gene count by computationally “contaminating” each cell type. For every pair of cell types, we randomly sampled and mis-labeled a fraction of cells (5% and 10%) and counted the number of expressed protein-coding (PC) and CRA genes at an expression cut-off of 0.03. We repeated this process 100 times to get a distribution of the number of genes expressed when cell types are incorrectly clustered (Supplementary Fig 11).

### P-value computation for sharing of gene sets

Expressed-gene sets were compared between all pairs of cell types using the Jaccard index. To test whether lineage-committed TECs share expressed genes more extensively among themselves than classical TECs, we computed the mean within-group Jaccard index for lineage-committed and classical TECs; the difference of these means served as our observed test statistic for comparison against an empirical null distribution. The null distribution was generated by relabeling cell types (all possible relabeling was enumerated exhaustively, amounting to 210 random groups) and computing the same test statistic. The empirical p value is the number of times the difference in means in groups of shuffled cell type labels is at least large as the observed value, divided by the 210, the total number of relabeled groups.

### Neighbor-voting analyses with MetaNeighbor

We ran MetaNeighbor [20] using the MetaNeighborUS function in the python pymetaneighbor implementation (version 0.1.0) (https://github.com/gillislab/pyMN). We used a down-sampling approach to account for differences in cluster size. Prior to each run of the MetaNeighborUS function, we downsampled each cluster to have the same number of cells as the smallest input cluster. We repeated this procedure 10,000 times and the final AUC scores were calculated as the means of the scores of each of the 10,000 runs.

### Single-cell ATAC-seq analysis of TECs

#### Reference genome

We used the same reference genomes for the GEX analyses and scATAC-seq analyses: GRCh38 for human samples and GRCm39 for mouse samples.

#### Read alignment and PCR duplicate removal

Cell barcodes were added to the FASTQ names field of the paired-end reads. Reads were mapped to the reference genome using BWA MEM (0.7.17) [66]. PCR duplicated were marked using samtools (1.17) [67].

#### Fragment file generation

SAM files were converted to fragment files retaining just the fragments with a minimum mapping quality of at least 30 and that are not PCR duplicates. As part of the fragments file generation, scATAC-seq barcodes were converted to the respective GEX barcode for the multiome datasets.

#### Cell-peak matrix generation

The cell-peak matrix was generated using MACS3 (3.0.1) [68] called via Signac (1.10.0) [69] using the default parameters.

#### Quality control

Quality control was performed in Signac (version 1.16.0) [69] using adaptive filters, with the goal of removing low-quality outlier cells (Supplementary Figs 3 to 5. For sample HT08, cells were removed if they had nCount_peaks below 3,000 and greater than 43,000, TSSE scores below 3, nucleosme signals above 3, a blacklist ratio above 0.01, and pct_reads_in_peaks below 35. For sample HT11, cells were removed if they had nCount_peaks below 700 and greater than 16,000, TSSE scores below 2.8, nucleosme signals above 3, a blacklist ratio above 0.01, and pct_reads_in_peaks below 30. Sample HT13 had overall poor QC metrics and was thus excluded from any downstream scATAC analysis (Supplementary Fig 5).

#### Motif Enrichment

We performed motif enrichment on the multiome dataset (TECs passing both scATAC-seq and snRNA-seq quality control) with chromVAR (version 1.24.0) [22] on the *Homo sapiens* motifs in the JASPAR2024 package (version 0.99.6) [70]. Motifs with the highest mean deviation score for each cell type were selected.

#### Joint scATAC and snRNA UMAP

We generated the joint UMAP for the multiome dataset using Seurat (version 5.3.1) [71] and Signac (version 1.16.0) [69]. We first generated an integrated latent semantic indexing (LSI) dimensionality reduction for the scATAC data using the FindIntegrationAnchors and IntegrateEmbeddings functions. We also generated an integrated count matrix for the snRNA data using the FindIntegrationAnchors and IntegrateData functions, and then calculated a PCA dimensionality reduction using the integrated counts. We used the FindMultiModalNeighbors function to construct a weighted nearest neighbor graph with the integrated LSI and PCA as inputs, and then used the RunUMAP function to generate the joint UMAP using this graph.

### Cell type lineage tracing using matched scRNA-seq and scATAC-seq data

#### Re-sequencing ATAC-seq libraries

Based on the QC and the sequencing depth, we re-sequenced the libraries using 100bp paired-end reads for HT08 and HT11 to gain sufficient coverage for lineage analysis, yielding 5,644 (HT08) and 2,677 (HT11).

#### Genome coverage analysis

The number and quality of mutations calls and the lineage trees built from these depend on the read coverage. To obtain coverage, we determined the mean read length as the weighted average of the read lengths weighted by the number of reads. For each cell type, we calculated the total number of fragments by summing the fragment count of the respective cells. We then multiplied the fragment count with the mean read length and divided by 3 billion to obtain the cell type coverage. The total sample coverage is the sum of all cell type coverages.

#### Pseudobulking of scATAC-seq reads

For lineage tracing, we only used cells that passed both snRNA-seq and scATAC-seq QC. Cell type assignments were based on the snRNA-seq data (as described above). Based on this assignment, we processed pooled scATAC-seq reads by reassigning single-cell read group identifiers (cell barcodes) in the BAM file using a barcode-to-cell type mapping with pysam (v) to create a single BAM file with reads grouped by cell type. The grouped BAM file was sorted and indexed using samtools (v).

#### Somatic variant calling with BCFtools

We used bcftools mpileup (v) to compute genotype likelihoods and allele-specific read depths across the cell-type–grouped BAM file, retaining only reads with base and mapping quality scores ≥ 20 (error probability ≤ 0.01). The resulting pileup BCF was filtered to keep only autosomal SNPs showing evidence for both alleles, removing homozygous reference and homozygous alternative sites. To exclude inherited variants, we used bcftools isec to subtract sites present in gnomAD (v4.1) with allele frequency > 0.001 (0.1%) across 76,215 whole genomes. MACS was applied to test whether each remaining site shows variation beyond that expected by sequencing error, retaining sites with QUAL *CLU*. This step removes sites with mostly homozygous reference alleles and few spurious variant reads but often retains sites that are mostly homozygous alternative with limited reference support. Because such sites are not informative for somatic mutation analysis, we implemented an additional pre-filter that reuses the genotype likelihoods to remove sites with *P* (*Data* ∣ all samples 0∕0) + *P* (*Data* ∣ all samples 1∕1) ≥ 0.001, leaving only loci with strong polymorphic signal for downstream testing.

#### Distinguishing somatic heterozygous sites from germline heterozygous sites

To distinguish somatic from germline heterozygous variants, we designed a likelihood ratio test (LRT) that compares a null model with a single shared allele frequency across all cell types to an alternative model allowing cell-type-specific allele frequencies. From the sites called in the previous step, a locus is considered for the LRT only if at least two cell types show germline homozygous signal (to reduce the influence of allelic dropouts) and at least one cell type shows evidence of the alternate allele, ensuring phylogenetically informative variation.

At one candidate site, each cell type *i* provides an observed number of alternate reads *k_i_* out of total reads *n_i_*. Cell types with no coverage (*n_i_* = 0) are excluded. The likelihood for a cell type under allele frequency *p* is modeled using the binomial distribution:

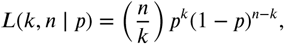

and the corresponding log-likelihood is

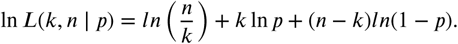

Under the null hypothesis *H*_0_, all *N* cell types are assumed to share a single global germline heterozygous allele frequency,

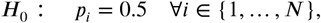

hence the null log-likelihood is given by

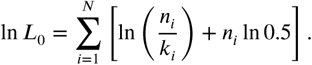

Under the alternative hypothesis *H*_1_, each cell type is allowed to have its own maximum likelihood estimate (MLE) of the allele frequency,

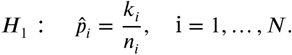

The corresponding log-likelihood under *H*_1_ is

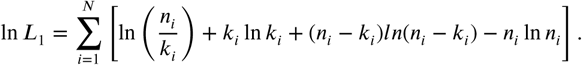

The likelihood ratio test statistic is then

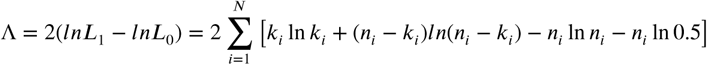

where the independence across cell types and computation in log space re-frames the LRT as a sum of the incremental improvements that allowing a free parameter provides over a fixed null heterozygous sampling per cell type.

Under asymptotic conditions, the likelihood ratio statistic Λ follows a *χ*^2^ distribution with degrees of freedom equal to the number of free parameters (i.e., the number of cell types) between the null and alternative models. However, because our data are limited and the asymptotic assumption may not hold, we instead estimate the null distribution of Λ empirically by Monte Carlo simulation.

Under the null hypothesis *H*_0_, new pseudo-datasets are generated by sampling alternate read counts *k*^(*s*)^ independently for each sample from a binomial distribution with parameters *n_i_* (the observed read depth) and *p*_0_ = 0.5 (the shared null frequency):

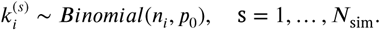

The log-likelihood ratio statistic is then computed for each replicate and the empirical *p*-value is computed as the proportion of simulated replicates whose likelihood ratio exceeds the observed statistic:

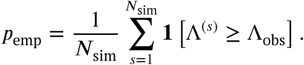

We used *N*_sim_ = 10, 000 simulations per site and the binomial draws preserve the original per-sample read depths to accurately reflect sampling variability. Sites with *p*_emp_ < 0.01 are considered to show significant deviation from the null expectation, indicating evidence of somatic heterozygosity.

#### Simulation to obtain empirical estimates of specificity and sensitivity of the LRT

To validate the final somatic mutation calls for each sample, we obtained the empirically observed read count totals per cell type per sample for each site in the final mutation calls. We then simulated 10 positive test cases, representing a true somatic heterozygous site, where mutations classified as germline (only evidence of the major allele in the empirical data) are simulated with all read counts as the major/ref allele (n ref reads; 0 alt reads) and somatic mutations are simulated as a heterozygote (alt reads drawn from binomial 0.5 distribution up to the total number of reads empirically observed per cell type). For the same site and to maintain positive/negative class balance, we simulated 10 negative test cases (meant to represent only a germline heterozygous site) where all germline and somatic mutations are simulated with alternative reads from the binomial(0.5). We applied the LRT described above to these 20 simulations per site and classified results as true positive, true negative, false positive, and false negative to obtain a sensitivity and specificity metric per mutation site.

#### Calling presence/absence of mutations for each cell type

After identifying candidate somatic heterozygous sites using the likelihood ratio test, we generate a discrete mutation matrix that distinguishes confidently mutated (evidence for the somatic heterozygous mutation) from non-mutated cell type populations (evidence only for the homozygous germline allele), while conservatively treating ambiguous low-depth cases as missing data.

For each candidate site, the reference (*r_i_*) and alternate (*a_i_*) read counts are extracted per cell type. The consensus call for cell type *i* is determined using a simple read-depth heuristic:

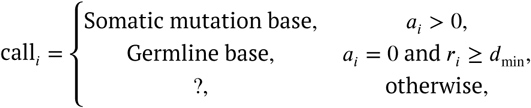

where *d*_min_ is a minimum read depth threshold (set to *d*_min_ = 2). Any cell type lacking alternate allele support and with fewer than two reference reads is marked as missing, since a single reference read is equally consistent with a reference or heterozygous genotype.

For both the LRT pre-filtering and final mutation calling, when homozygous samples exist for both alleles, the majority allele is treated as the germline state, and the minor allele is interpreted as evidence of a somatic heterozygous mutation, assuming homozygosity for the minor allele arises from low coverage or allelic dropout.

#### Phylogenetic inference

We inferred phylogenetic trees of cell type lineages using both Bayesian (MrBayes v3.2.7a) and maximum likelihood (RAxML-NG v1.2.2) methods. For each sample, we concatenated our high confidence somatic SNP sites into a sequence alignment of consensus genotypes for each cell type (either in FASTA or NEXUS file format). We then specified the HKY nucleotide substitution model for both methods and used the default settings for Bayesian priors and parameter initializations. For Bayesian inference, we ensured convergence to the posterior distribution by standard criteria (effective sample size and average standard deviation of split frequencies) and drew 10,000 samples to summarize phylogenetic trees and parameters. For maximum likelihood inference, we first inferred a single best tree and then ran 1000 bootstrap tree estimates to overlay confidence values. Results for sample HT11 are shown in Figure 2, for sample HT08 in Supplementary Fig 9

#### Lineage analysis of Gamble et. al. dataset

The fastq files were processed as described in *Single-cell ATAC-seq analysis of TECs*. The cell annotations were obtained from the file Source Data Extended Data Fig.1 (Mimetic.Cluster column) in [19].

### SCENIC+ analysis of TECs

We used the SCENIC+ pipeline [30] (version 1.0a1) to identify regulons associated with each TEC subtype in the human multiome dataset (this study) and the mouse multiome dataset from Givony *et al.* [11]. The following paragraphs describe the pre-processing steps and parameters for setting up the pipeline.

#### pycisTopic analysis

We used pycisTopic [30] (version 2.0a0) to generate pseudobulk fragments files for each cell type and called peaks using MACS [68] (version 2.2.9.1). We inferred consensus peaks twice, first for each pseudobulk fragments file and then for all of the peaks together, generating a consensus bed file to be used both for subsequent topic modeling and later in the SCENIC+ pipeline. Using the consensus bed file, we ran Latent Dirichlet Allocation topic models for the multiome data with n_topics = [2, 5, 10, 15, 20, 25, 30, 35, 40, 45, 50]. We selected the number of topics to use in the downstream analysis with the evaluate_models function.

#### Generation of custom databases for pycisTarget

We followed the tutorial at https://scenicplus.readthedocs.io/en/ latest/human_cerebellum_ctx_db.html to generate custom databases of ranked and scored motifs for the consensus peaks generated by pycisTopic.

#### SCENIC+ Motif Analysis and Regulon Identification

We implemented the SCENIC+ Snakemake pipeline outlined in the documentation located at https://scenicplus.readthedocs.io/en/latest/index.html. All parameters were kept at default but for min_target_genes, which was set to 5. Motif enrichment was run via pycisTarget [30] (version 1.0a2) using the custom motif ranking and scoring databases described above. SCENIC+ was then used to link TF’s, target genes, and genomic regions, yielding TF-gene, TF-region, and region-gene modules. The grn_inference function was used to aggregate the modules into regulons. The AUCell function was used to generate gene-based and region-based regulon AUC scores for each cell. We then assigned a cell type to each regulon, selecting the cell type with the highest gene-based AUC score.

Regulons were identified as activators or repressors based on their TF-to-gene and region-to-gene relationships, as indicated by the suffix in the regulon name provided by SCENIC+. A positive TF-to-gene relationship indicates that when the regulon-driving TF is expressed, the target genes are also expressed, while a negative TF-to-gene relationship indicates that when the regulon-driving TF is expressed, the target genes are not expressed. A positive region-to-gene relationship indicates that the chromatin regions linked to the target genes are open when the target genes are expressed. Regulons with a positive TF-to-gene relationship and a positive region-to-gene relationship (denoted by the (+/+) suffix) were determined to be activators. Regulons with a negative TF-to-gene relationship and a positive region-to-gene relationship (denoted by the (-/+) suffix) were determined to be repressors.

#### Over-Representation Analysis

We performed over-representation analysis on the target genes of each of the regulons identified by the SCENIC+ analysis using GSEAPY (version 1.1.9) [72]. We accessed the Enrichr API [73] from within the GSEAPY package and performed over-representation analysis with the enrichr function, comparing the representation of input gene sets in the regulon target genes to their representation in a set of background genes. We used a subset of gene sets from the GO [74, 75], KEGG [76] and MsigDB [77–80] databases as well as gene sets from the Tabula Sapiens [81, 82] and Tabula Muris [83] databases (Supplementary Table 10). The full set of genes in the multiome snRNA-seq objects were set as background genes. After removing pathways that were duplicated across gene sets (keeping the first instance of a given pathway and removing only the duplicates), we identified significantly over-represented pathways using a threshold of 0.05 for the adjusted p-values output by GSEAPY, and identified the top over-represented pathways for each regulon based on the Combined.Score output.

We selected the pathways shown in Figure 4F and Figure 6F with the goal of maximizing the number of displayed cell-type regulons. For cell types with more than 2 assigned regulons, we selected the pathway for each regulon with the largest Combined.Score value to generate a set of top regulon-pathway pairs. We chose to display at most 5 regulon-pathway pairs for each lineage-committed TEC subtype, so if the lineage-committed TEC subtype had more than 5 assigned regulons, the 5 regulon-pathway pairs with the greatest Combined.Scores were chosen to display. If a cell type had 2 assigned regulons, the top 2 pathways for each regulon were selected to display based on their Combined.Scores, for a total of 4 displayed regulon-pathway pairs. If a cell type only had 1 assigned regulon, then the top 3 pathways for each regulon were selected to be displayed, for a total of 3 displayed regulon-pathway pairs.

## Author contributions

Theoretical and computational analysis: RP, SRC, SJS, DP, AJK, AG, HVM. Experimental data generation and processing: YL, DN, ML, SC, JT, TC, CR, HVM; Tissue acquisition and processing: HP, RM; Writing - original draft: RP, SRC, YL, HVM; Writing -review & editing: all authors; Conceptualization, funding acquisition: HVM; Supervision: AS, JP, HVM.

## Code availability

Custom analysis code was written in either R (version ≥ 4.2.0) or python (version ≥ 3.11). All analysis code will be freely available on GitHub: https://github.com/meyer-lab-cshl/intecgrated_2026.

## Data availability

FASTQ files and processed data objects (seurat, anndata) as well as additional metadata will be available from dbGAP upon publication (access controlled for raw sequencing files). Data to reproduce figures are provided in the github repository above. The integrated human and mouse datasets can be viewed via a Shiny app located at http://intecgrated.cshl.edu/.

## Acknowledgments

We thank Jacqueline Bott and the staff at CTSL for study coordination and administrative support.

## Funding

The research was supported by the Simons Center for Quantitative Biology at Cold Spring Harbor Laboratory; the Cold Spring Harbor Laboratory and Northwell Health Affiliation; National Science Foundation Graduate Research Fellowship (to SJS); Starr Centennial Scholarship (to SJS); US National Institutes of Health (NIH) National Institute of General Medical Sciences Grant R35-GM127070 (to AS); US National Institutes of Health Grant S10OD028632-01 and 1R01AI167862 (to HVM); and the Simons Pivot Fellowship (to HVM). This work was performed with assistance from the Cold Spring Harbor Laboratory Shared Resources, including the Flow Cytometry, Next Generation Sequencing and Single Cell Shared Resources, which are supported by the Cancer Center Support Grant 5P30CA045508. This work was also supported by the CSHL Microscopy Shared Resource and National Institutes of Health Shared Instrument Grant 1S10OD034372-01, which funded the purchase of the Zeiss LSM 980 with Airyscan 2. It was further supported by the Center for Biospecimen Research and Development which is partially supported by NYU Langone’s Laura and Isaac Perlmutter Cancer Center support grant P30CA016087. The funders had no role in the study design or decision to publish.

During the preparation of this work, the authors used *Cursor* in *auto* mode to tighten language and check for inconsistencies. The authors reviewed and edited the output as needed and take full responsibility for the content of the published article.

