## Supplementary Figures and Tables for "Gene regulatory logic underlying the differentiation of mixed lineages in human thymic epithelial cells"

**Supplementary Table 1. CELLxGENE tissue type annotations.** Data are in the corresponding spreadsheet. *cellxgene_tissue*: tissue type label in CELLxGENE; *cellxgene_tissue_general*: higher-order ontology of the tissue label in CELLxGENE; *tissue_curated*: manually and ontology-based curated tissue type. Tissue types with QC_fail entries were removed from downstream analyses. *cell_count*: total number of cells assigned to the tissue.

**Supplementary Table 2. CELLxGENE cell type annotations.** Data are in the corresponding spreadsheet. *celltype_cellontology*: cell type label in CELLxGENE; *celltype_curated*: manually and ontology-based curated cell type. Cell types with QC_fail entries were removed from downstream analyses.

**Supplementary Table 3. Cell type-restricted antigens and tau scores for module scoring: human.** Data are in the corresponding spreadsheet. *Gene*: gene name; *Tau*: estimated tau value for cell-type specificity; *Cell type*: curated CELLxGENE cell type to which the cell-type-restricted antigen is assigned.

**Supplementary Table 4. Donor characteristics.** Data in corresponding spreadsheet. *study*: name of study used throughout manuscript; *sample*: sample identifier; *individual*: individual identifier; *sex*: male/female; *age*: age at time of surgery (d: days, m: months, y:years); *reason for surgery*: reason for surgery as indicated at collection

**Supplementary Table 5. Accession numbers and download links for public single-cell and single-nuclear RNA sequencing dataset of human and mouse thymus.** Dataset: identifier (first author in manuscript) used throughout this study.

| Dataset | Species | Download location | Accession |
|---|---|---|---|
| michelson | mouse | https://www.ncbi.nlm.nih.gov/geo/query/acc.cgi?acc=GSE194252 | GSE194252 |
| givony | mouse | https://www.ncbi.nlm.nih.gov/geo/query/acc.cgi?acc=GSE236667 | GSE236667 |
| raggazzini | human | https://www.ncbi.nlm.nih.gov/geo/query/acc.cgi?acc=GSE220830 | GSE220830 |
| bautista | human | https://www.ncbi.nlm.nih.gov/geo/query/acc.cgi?acc=GSE147520 | GSE147520 |
| park | human | https://www.ebi.ac.uk/biostudies/arrayexpress/studies/E-MTAB-8581 | E-MTAB-8581 |
| gale | mouse | https://www.ebi.ac.uk/biostudies/arrayexpress/studies/E-MTAB-8560 | E-MTAB-8560 |
| kousa | mouse | https://www.ncbi.nlm.nih.gov/geo/query/acc.cgi?acc=GSE240020 | GSE240020 |
| huisman | human | https://www.ncbi.nlm.nih.gov/geo/query/acc.cgi?acc=GSE262749 | GSE262749 |
| campinoti | human | https://www.ncbi.nlm.nih.gov/geo/query/acc.cgi?acc=GSE159745 | GSE159745 |
| gamble | mouse | https://www.ncbi.nlm.nih.gov/geo/query/acc.cgi?acc=GSM8447493 | GSM8447493 |

**Supplementary Table 6. Metadata for each of the public single-cell and single-nuclear RNA sequencing dataset of human and mouse thymus.** Data in corresponding spreadsheet.

**Supplementary Table 7. Parameters used for each dataset for reference genome indexing and read alignment with STARsolo.** Data in corresponding spreadsheet.

**Supplementary Table 8. TEC annotations based on marker genes: human.** Data are in the corresponding spreadsheet. *Cell type*: annotated cell type; *Marker*: gene name; *Source*: original study (by first author).

**Supplementary Table 9. TEC annotations based on marker genes: mouse.** Data are in the corresponding spreadsheet. *Cell type*: annotated cell type; *Marker*: gene name.

**Supplementary Table 10. Gene sets used for over-representation analysis.** *Gene Set*: gene set ID in GSEAPY.

| Gene Set | Species |
|---|---|
| GO_Biological_Process_2025 | Human, Mouse |
| GO_Cellular_Component_2025 | Human, Mouse |
| KEGG_2021_Human | Human |
| MSigDB_Hallmark_2020 | Human, Mouse |
| Tabula_Sapiens | Human |
| KEGG_2019_Mouse | Mouse |
| Tabula_Muris | Mouse |

### Figures

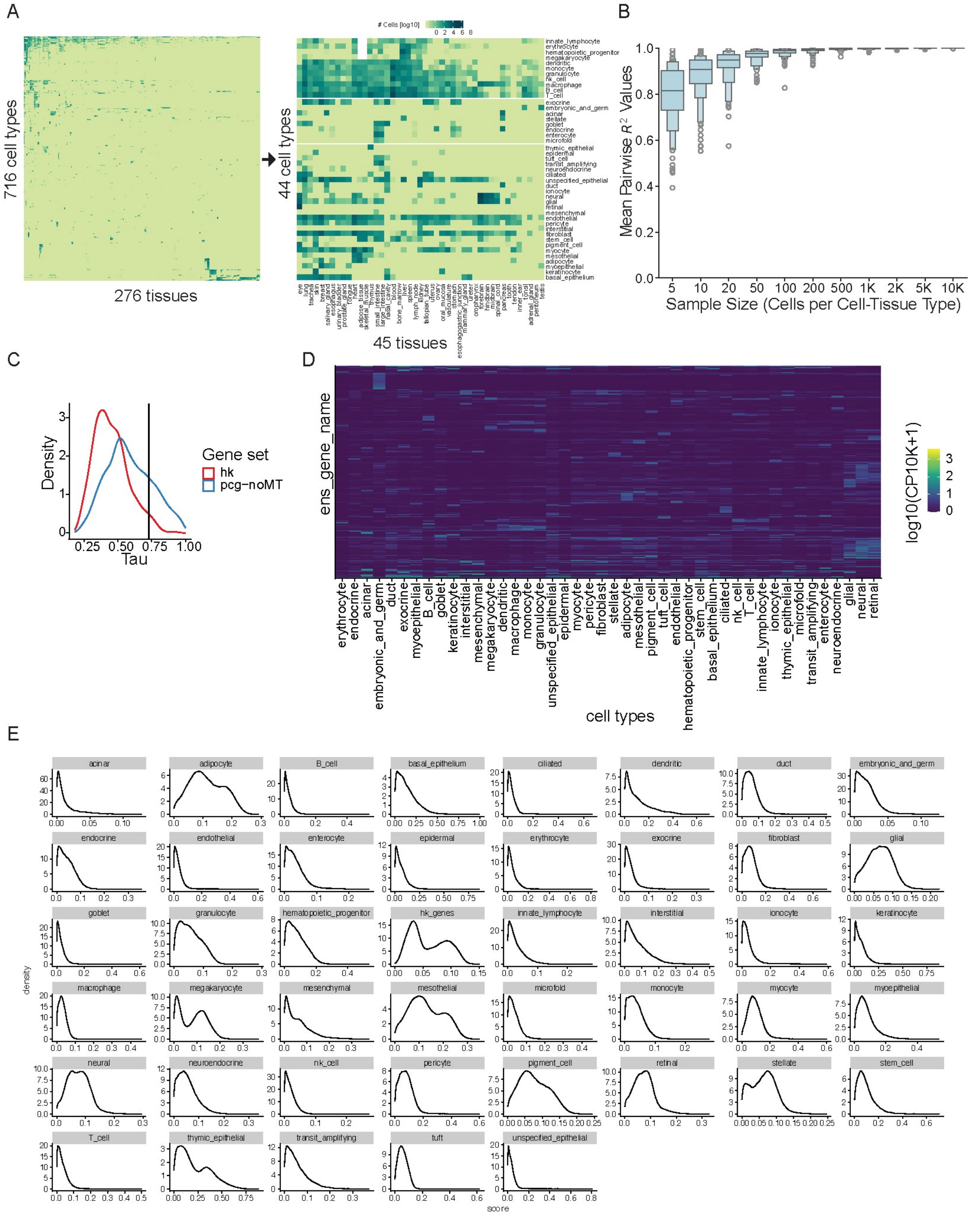


**Supplementary Figure 1. CELLxGENE and CRA scores. (A)** Cell numbers by cell type and tissue in the full CELLxGENE database [1] (left) and consolidated cell types and tissues (right). Only the lower block of cell types passed the signature score expression cut-off ( see methods); those are the scores shown in Figure 1A. **(B)** Pairwise Pearson's $R^2$ for 10 independent down-sampled datasets. X-axis indicates the number of cells per cell type per tissue. **(C)** Density distribution of estimated Tau values in house-keeping genes (HK) and protein-coding, non-mitochondrial genes (pcg-noMT). **(D)** Heatmap of log-transformed, normalized to 10k reads of genes with tau > 0.73, considered cell-type-restricted antigens (CRAs).

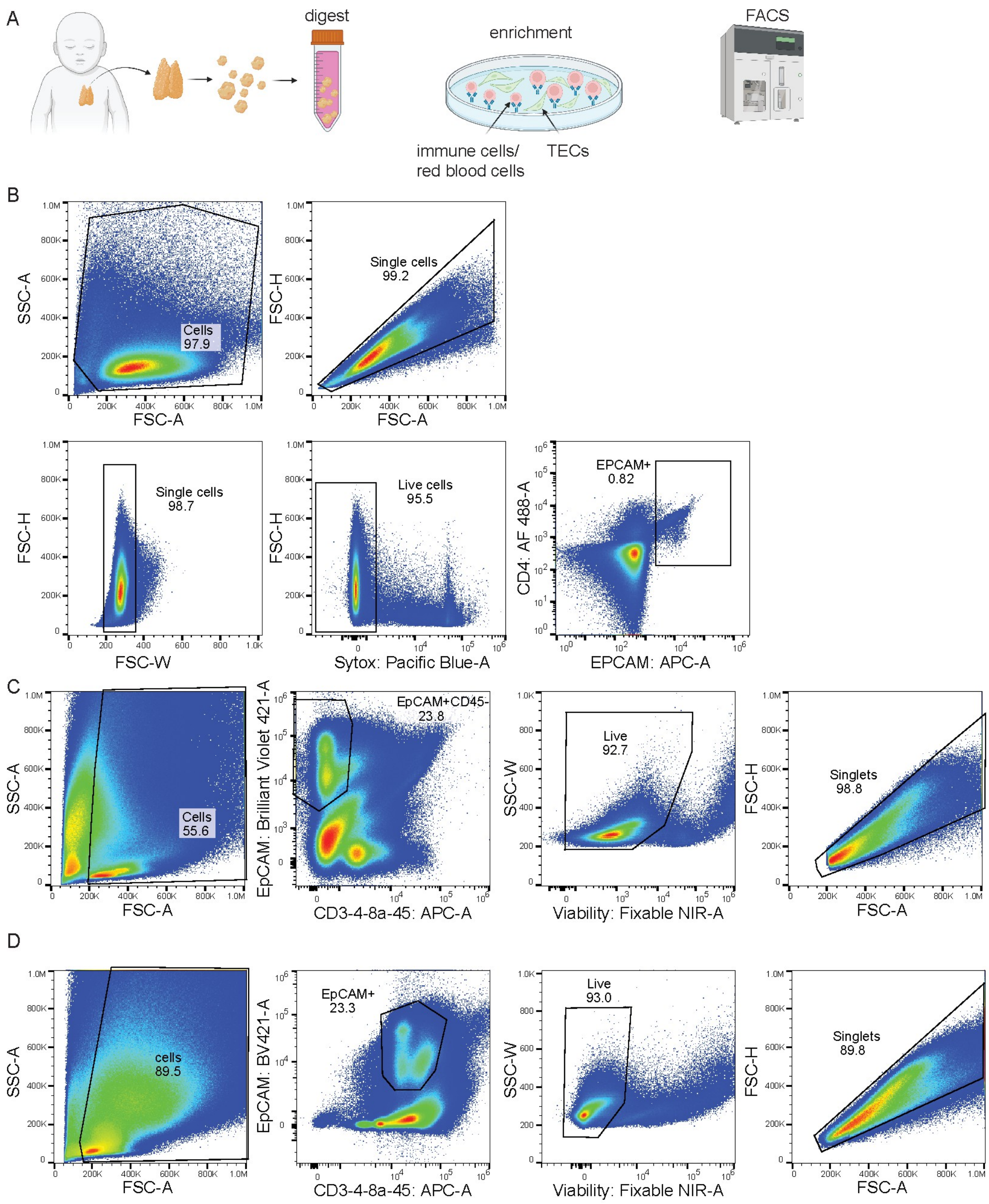


**Supplementary Figure 2. Human Thymus Processing and sorting for single-cell sequencing. (A)** Fresh human thymus tissue (donor characteristics in Supplementary Table 4) was delivered to the lab within 4 hours post-surgery. Up to 2 grams of the tissue was used for thymic epithelial cell isolation. The tissue was dissected with a scissor into progressively finer pieces and shaken to remove the thymocytes. After thymocyte release, the residual tissue was digested with an enzyme cocktail containing Collagenase IV, Dispase I, and DNase I, to release the epithelial cells from the tissue matrix. An enrichment step was performed using the panning technique whereby unwanted cells were removed through adherence to a petri dish via antibody-mediated binding. The TEC-enriched cell suspension was then stained and TECs sorted by FACS. **(B)** FACS strategy for sorting out TECs for multi-ome sequencing of the HT8 sample. A general cell gate was used first to remove obvious doublets and debris. Two rounds of singlet gating and dead cell exclusion was performed before gating on EpCAM+ cells for sorting. **(C)** FACS strategy for sorting out TECs for multi-ome sequencing of the HT11 and HT12 samples (representatively shown for HT11), a general cell gate was used first to remove debris and dead cells, followed by gating on EpCAM+CD45- cells. Dead cells were removed next, and lastly the singlet gate was used for sorting. **(D)** FACS strategy for sorting out TECs for PARSE sequencing (analogous to (C)).

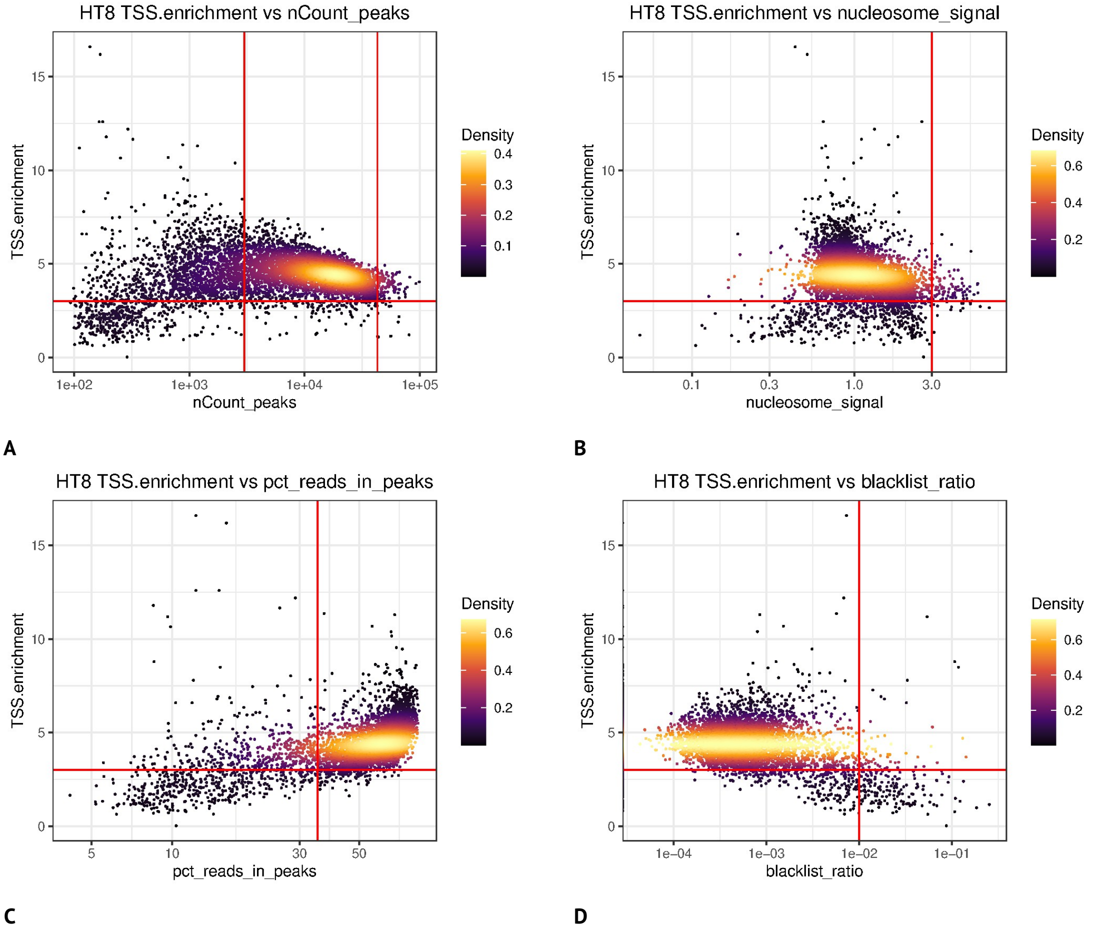


**Supplementary Figure 3. scATAC-seq QC for sample HT08 (A)** TSS.enrichment vs nCount_peaks. Using adaptive filters, cells passed QC if 3,000 < nCount_peaks < 43,000 and TSS.enrichment > 3. **(B)** TSS.enrichment vs nucleosome_signal. Using adaptive filters, cells passed QC if nucleosome_signal < 3 and TSS.enrichment > 3. **(C)** TSS.enrichment vs pct_reads_in_peaks. Using adaptive filters, cells passed QC if pct_reads_in_peaks > 35 and TSS.enrichment > 3. **(D)** TSS.enrichment vs blacklist_ratio. A standard cutoff of blacklist_ratio < 0.01 was used in addition to the adaptive filter of TSS.enrichment > 3. Cells passed QC if blacklist_ratio < 0.01 and TSS.enrichment > 3.

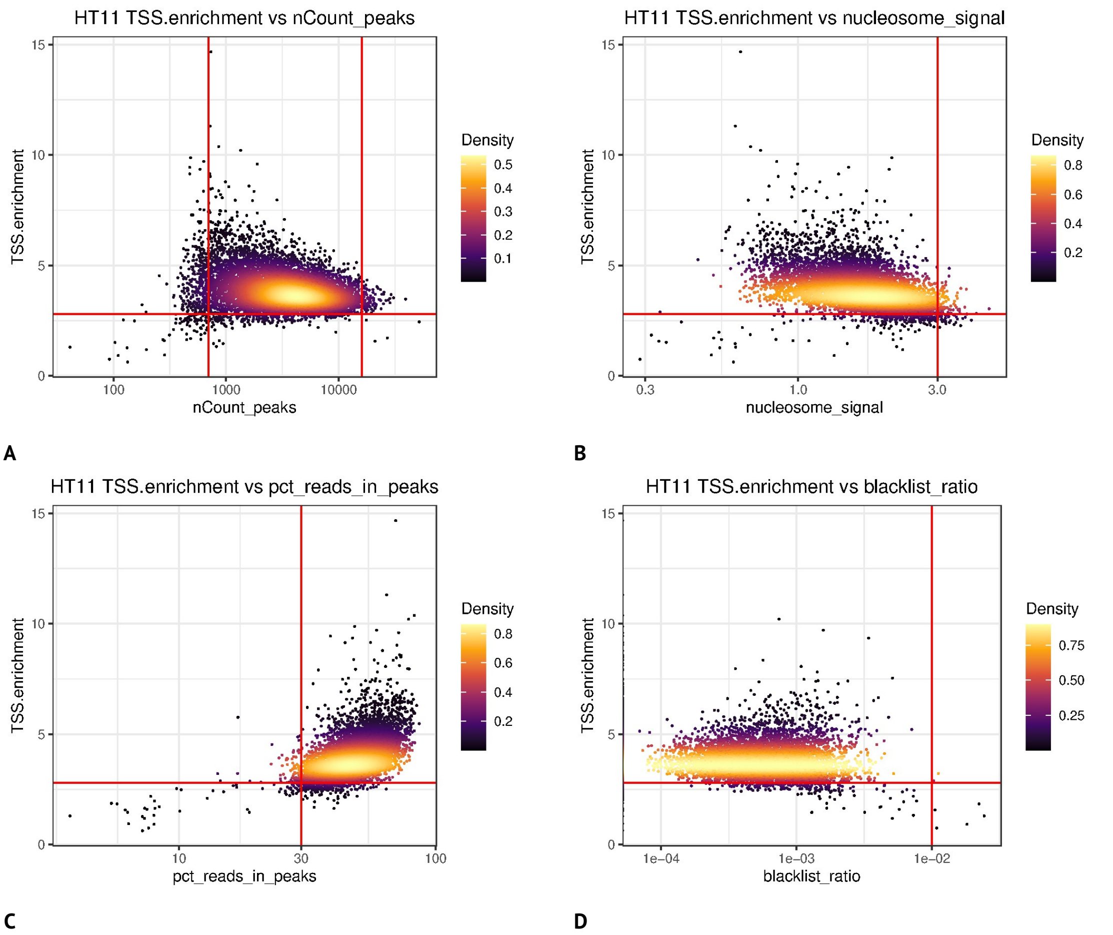


**Supplementary Figure 4. scATAC-seq QC for sample HT11 (A)** TSS.enrichment vs nCount_peaks. Using adaptive filters, cells passed QC if 700 < nCount_peaks < 16,000 and TSS.enrichment > 2.8. **(B)** TSS.enrichment vs nucleosome_signal. Using adaptive filters, cells passed QC if nucleosome_signal < 3 and TSS.enrichment > 2.8. **(C)** TSS.enrichment vs pct_reads_in_peaks. Using adaptive filters, cells passed QC if pct_reads_in_peaks > 30 and TSS.enrichment > 2.8. **(D)** TSS.enrichment vs blacklist_ratio. A standard cutoff of blacklist_ratio < 0.01 was used in addition to the adaptive filter of TSS.enrichment > 2.8. Cells passed QC if they had blacklist_ratio < 0.01 and TSS.enrichment > 2.8.

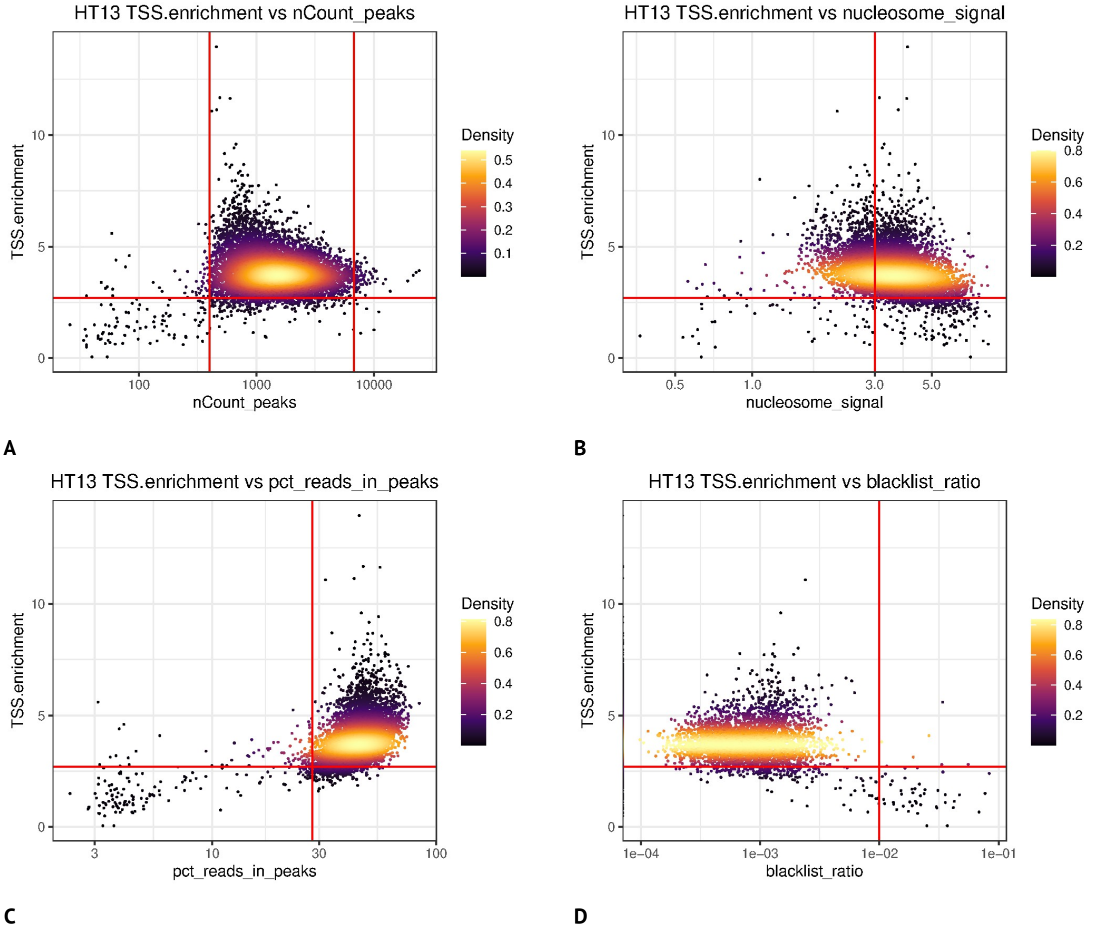


**Supplementary Figure 5. scATAC-seq QC for sample HT13, which was discarded due to poor library quality. (A)** TSS.enrichment vs nCount_peaks. An adaptive filter would have kept cells with TSS.enrichment > 2.7 and 400 < nCount_peaks < 6,750. The low values of nCount_peaks indicate a poor library quality. **(B)** TSS.enrichment vs nucleosome_signal. An adaptive filter would have set a threshold at nucleosome_signal = 6, an indication of the poor quality of the library. Removing cells with a nucleosome_signal > 3, as was done in samples HT8 and HT11, would also result in very few retained cells due to the poor quality of the library. **(C)** TSS.enrichment vs pct_reads_in_peaks. **(D)** TSS.enrichment vs blacklist_ratio.

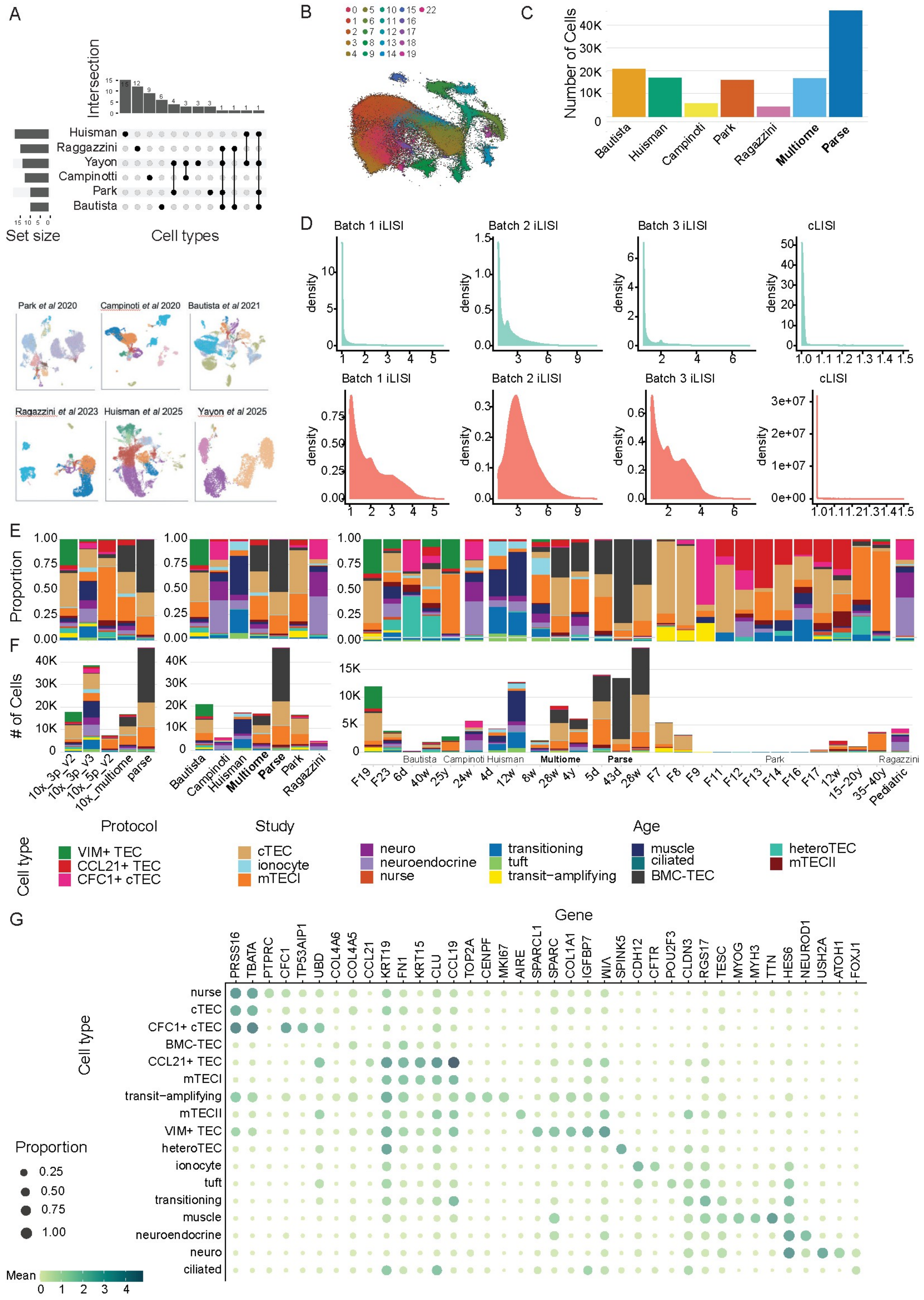

**Supplementary Figure 6. Data integration, clustering and annotation. (A)** Individual study cell embeddings (lower part) and size and intersection of cell types used in 6 prior human thymus single-cell datasets [2–7]. **(B)** Leiden clustering on *scVI* embedding of QC-ed, integrated and batch-corrected single-cell data from this study and prior studies in **(A)**. **(C)** Number of cells contributed by each study. **(D)** Degree of mixing during batch correction and dataset integration measured as the local inverse Simpson's index (LISI). The integration LISI (iLISI) measures the effective number of datasets within a neighborhood and cell-type LISI (cLISI) evaluates the accuracy of cell-type assignment. Mixing was assessed on sequencing protocol (Batch 1), donor (Batch 2) and study (Batch 3). Blue curves indicate LISI before integration, red after integration. Distribution of the proportion **(E)** and absolute number **(F)** of cells in each cluster by sequencing protocol, study and age. **(G)** Dotplot of mean normalized expression of marker genes for each cell type.

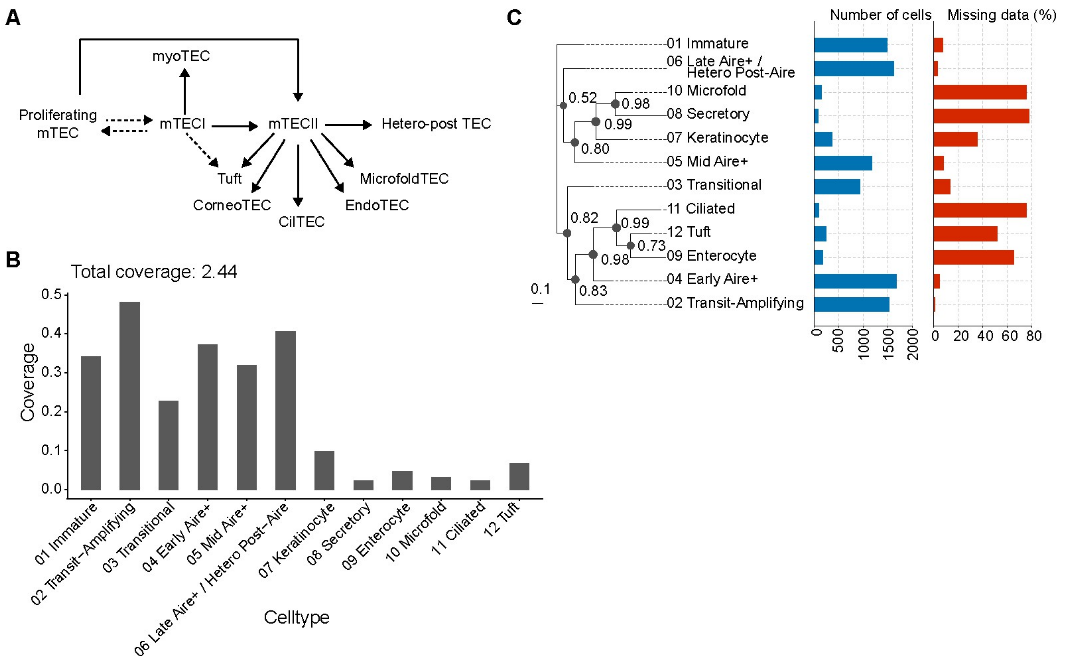


**Supplementary Figure 7. Murine TEC lineage tracing using somatic variant calling. (A)** Murine TEC lineage relationships as inferred by experimental lineage tracing by Givony *et al*; diagram based on extended Figure 9 in [8]. **(B)** Cell type-specific read coverage plots (x-axis) and total coverage for multi-ome data from Gamble *et al.* [9]. **(C)** Murine TEC lineage tracing based on somatic variant calling and phylogenetic inference on multi-ome data from Gamble *et al.* [9]. Numbers at branch points indicate posterior probabilities.

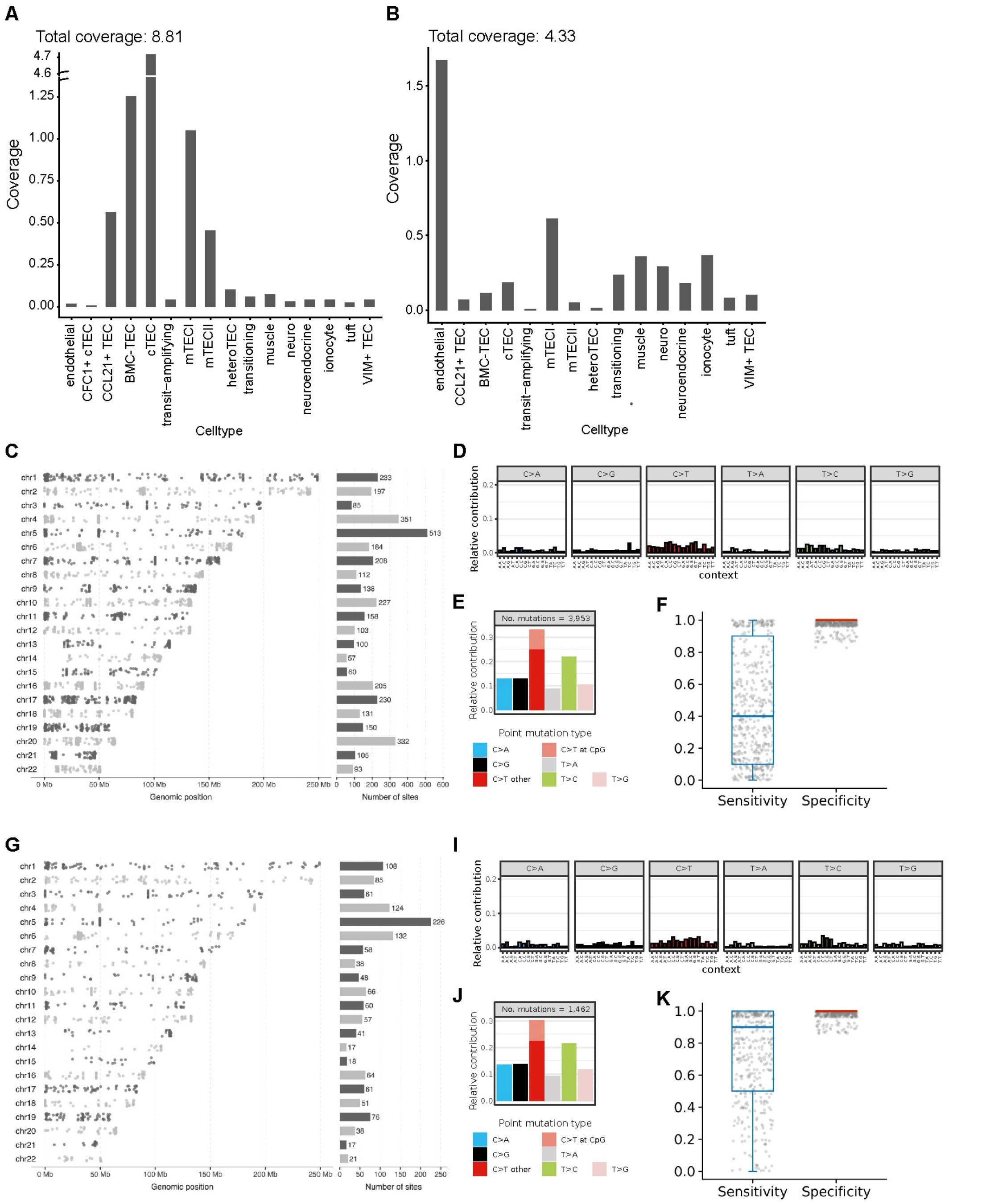


**Supplementary Figure 8. Somatic variant calling from scATAC-seq reads aggregated by cell type and sample.** **(A)** and **(B)** Cell type-specific read coverage plots (x-axis) and total coverage for samples HT08 and HT11, respectively. **(C)** Post-QC somatic mutation calls in sample HT11 are evenly distributed across chromosomes. **(D)** and **(E)** Types of mutations in the post-QC somatic mutation calls in sample HT11, as analyzed using the R package *MutationalPatterns* (v3.19.1). Minor elevation of C>T mutations supported by [10–12] as a common somatic mutation type due to DNA damage / improper DNA repair and replication errors at CpG sites. **(F)** Empirical sensitivity and specificity estimates from the LRT on simulated mutations. The high specificity lends confidence in the somatic mutation calls as true mutation sites. **(G)** - **(K)**: results as in **(C)** - **(F)** for sample HT08.

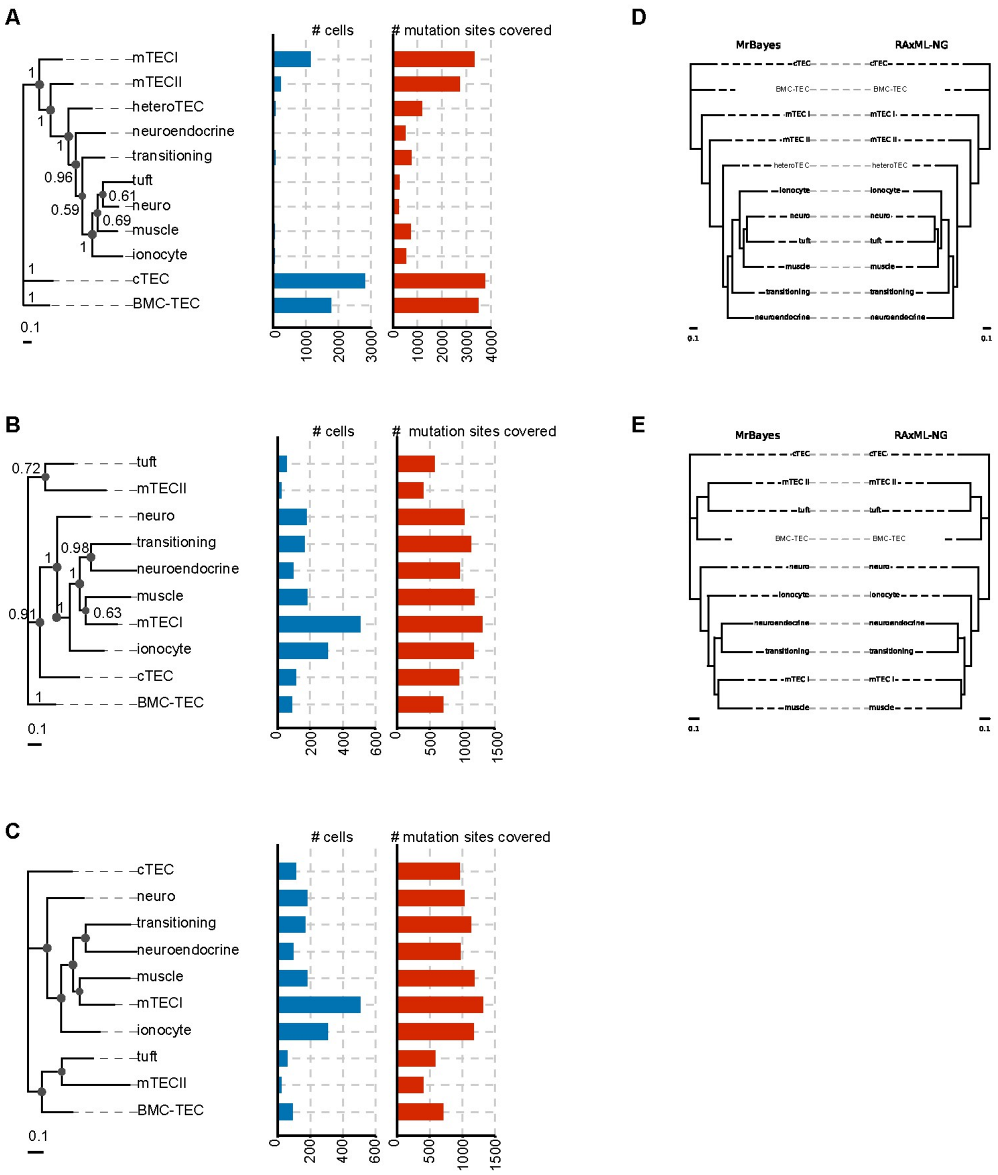


**Supplementary Figure 9. Human TEC lineage tracing using somatic variant calling. (A)** and **(B)** Unrooted phylogenetic tree for lineage-committed cell types with <5% missing data (left) for samples HT11 and HT08. Numbers at branch points indicate posterior probabilities. **(C)** Phylogenetic tree rooted on cTECs for lineage-committed cell types with <5% missing data (left) for sample HT08. Posterior probabilities as in (B). **(D)** and **(E)** Concordance of Bayesian tree (mrbayes) and maximum likelihood tree (raxml-ng) for samples HT11 and HT08.

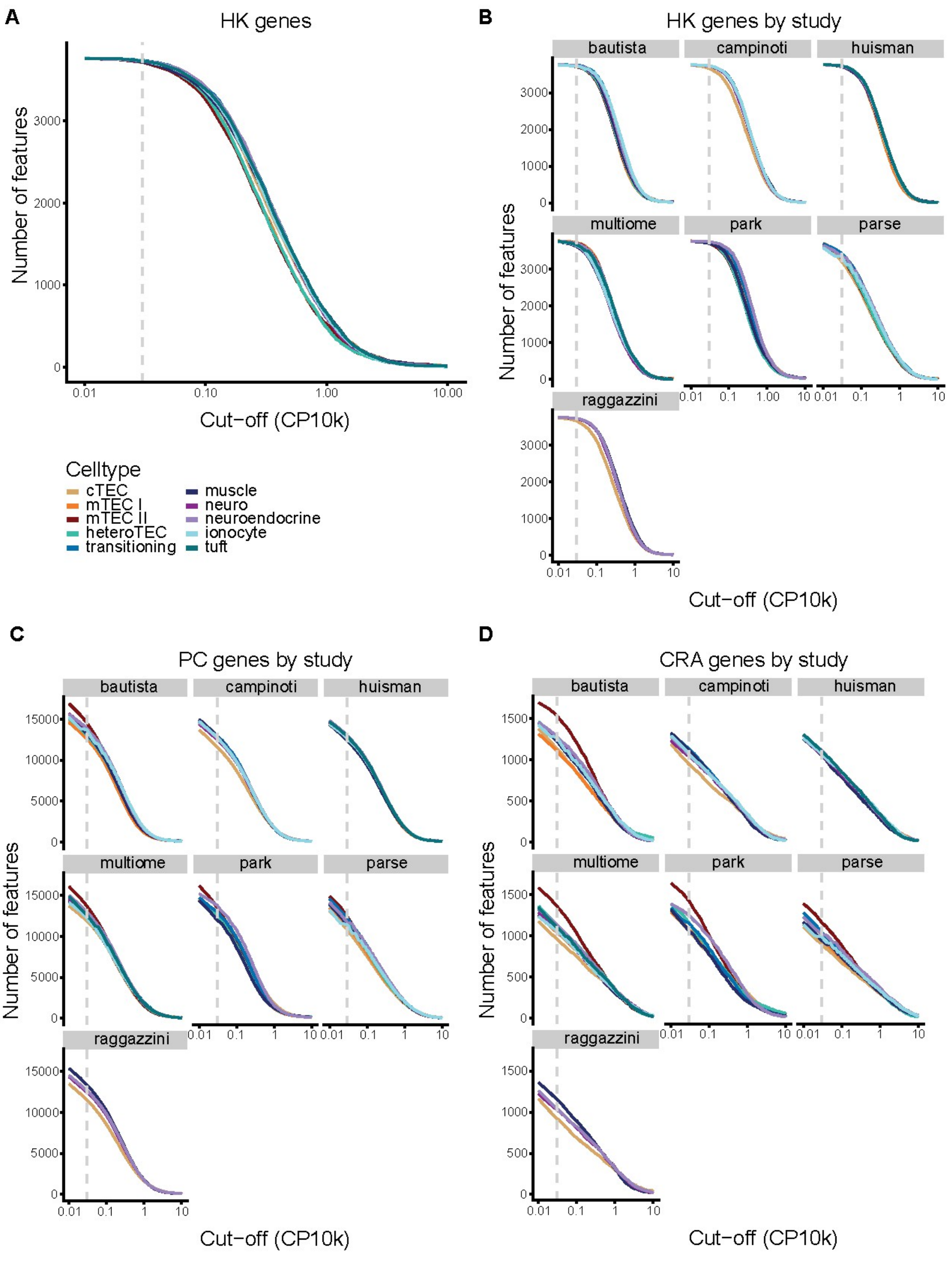


**Supplementary Figure 10.** Number of genes expressed at a range of cut-off values. **(A)** House-keeping (HK) genes. **(B)** HK genes split by study. **(C)** Protein-coding (PC) genes split by study. **(D)** Cell-type-restricted antigen (CRA) genes split by study.

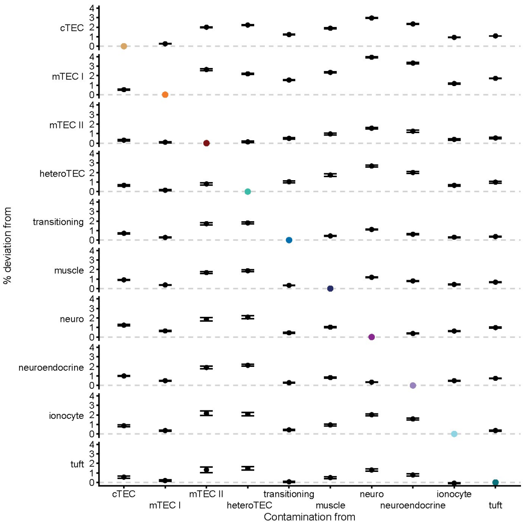


**A.** PC genes at 5% contamination

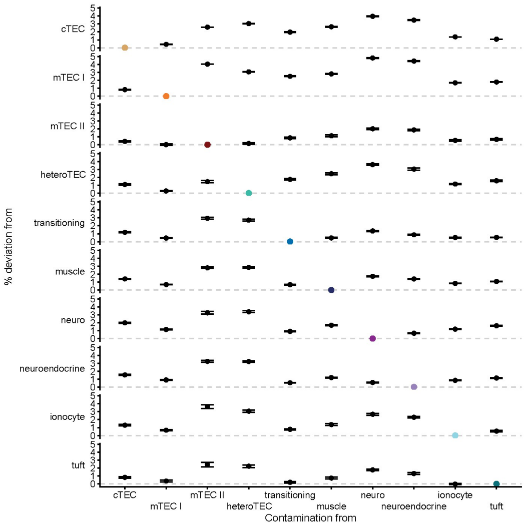


**B.** PC genes at 10% contamination

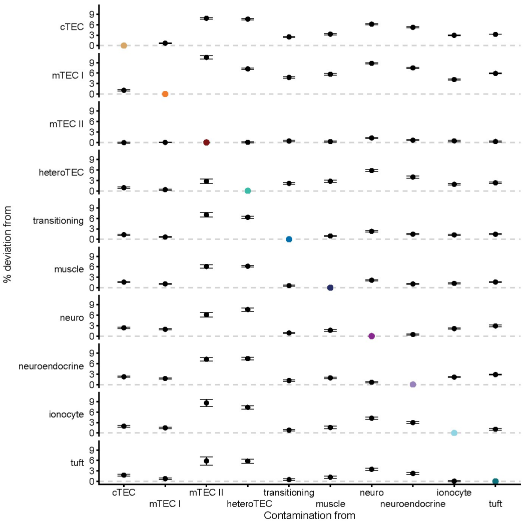


**C.** CRA genes at 5% contamination

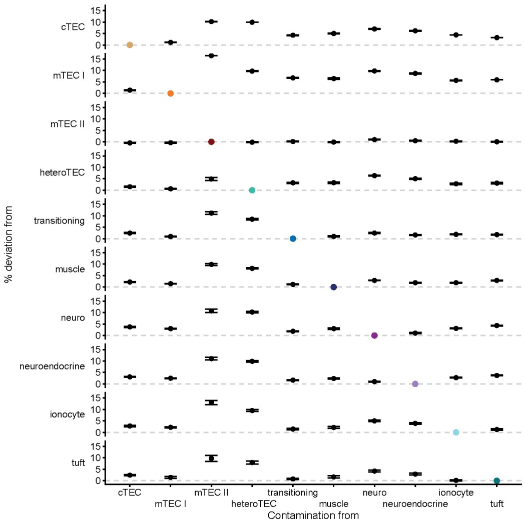


**D.** CRA genes at 10% contamination

**Supplementary Figure 11. Contamination analysis at an expression cut-off of 0.03.** For every pair, the distribution of the number of the genes when contaminated differs from the number of genes expressed by that cell type (gray line in each row). For instance, the least deviation was when 5% ionocytes were contaminated into tuft TD-TECs, while the deviation was more prominent at 10% contamination. On the other extreme, when cells were contaminated into mTECII, they reduced the number of genes expressed likely due to the low level of expression of promiscuously expressed genes.

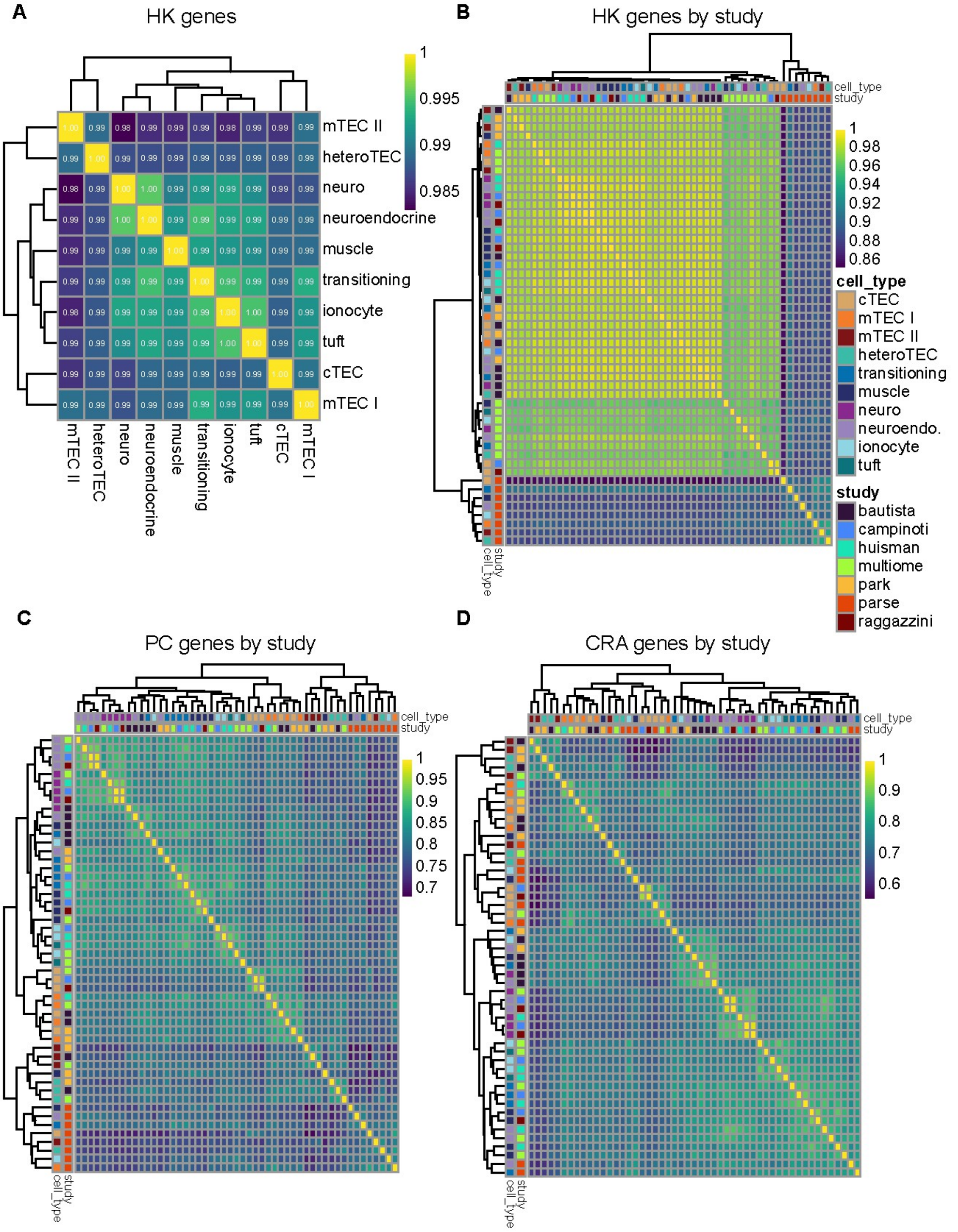


**Supplementary Figure 12.** Jaccard index of shared genes at a cut-off of 0.03. **(A)** House-keeping (HK) genes. **(B)** HK genes split by study. **(C)** Protein-coding (PC) genes split by study. **(D)** Cell-type-restricted antigen (CRA) genes split by study.

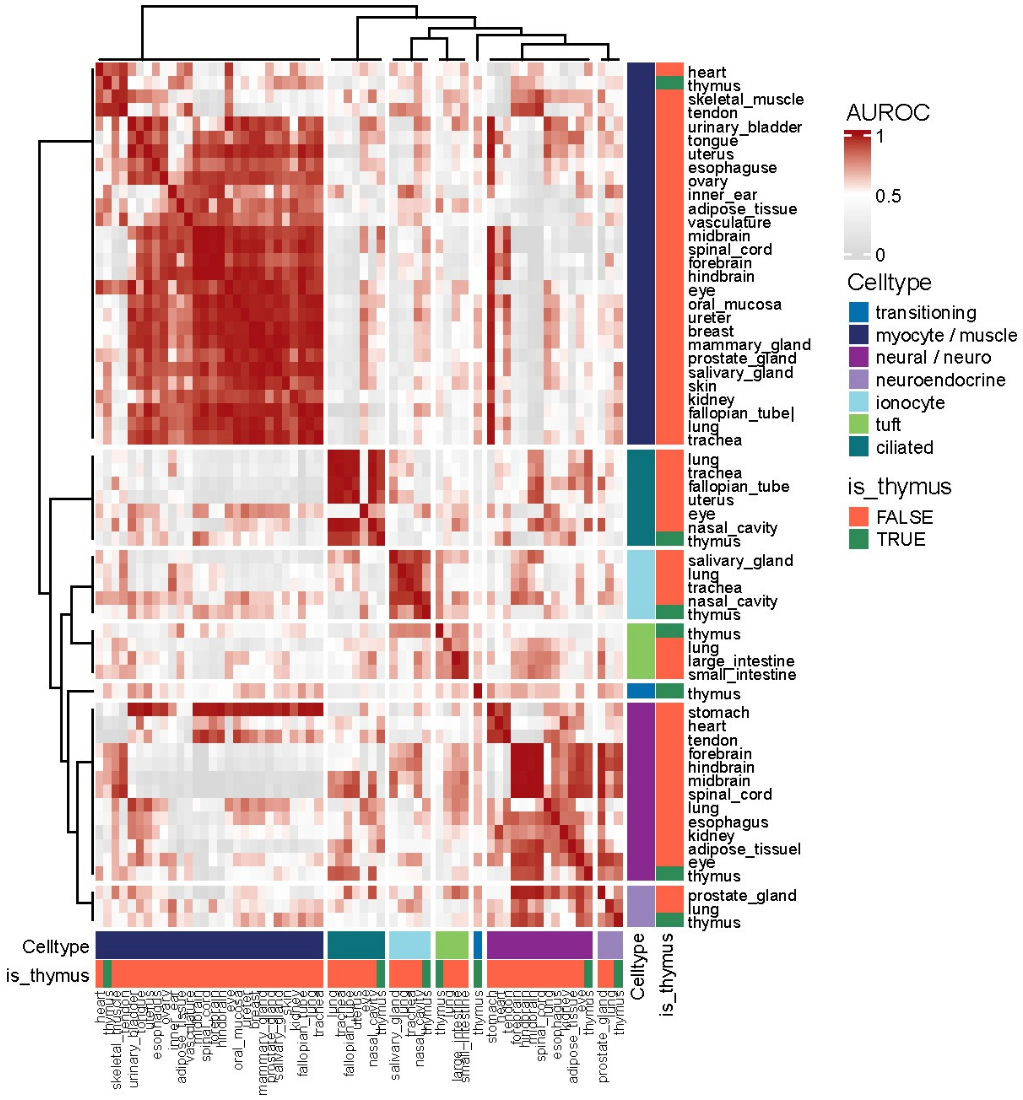


**Supplementary Figure 13.** Neighbor-voting meta-analyses (using MetaNeighbor) showing the similarity between every pair of cell types in the peripheral tissues and the thymus.

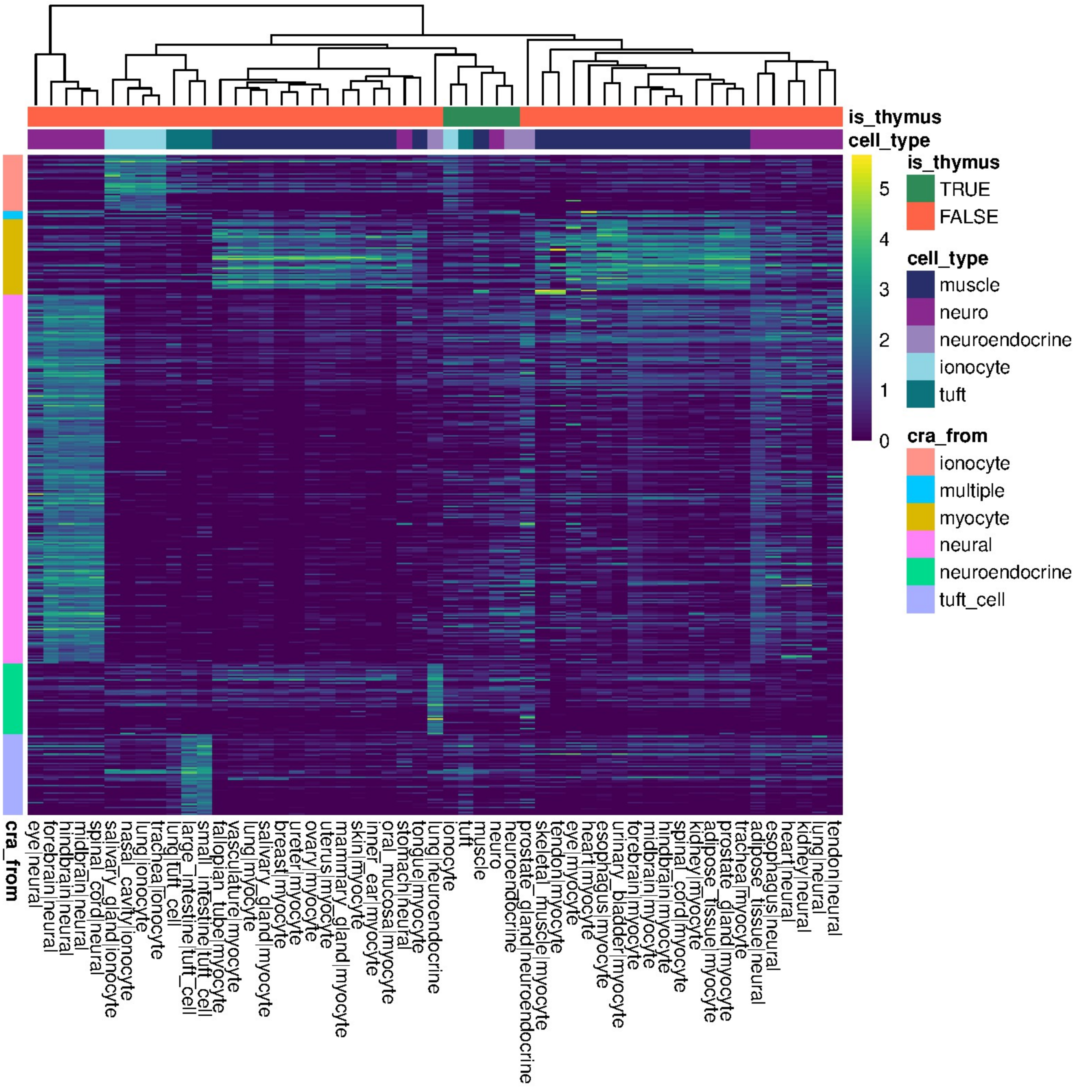


**Supplementary Figure 14.** CRA gene expression in thymus and peripheral tissue cell types.

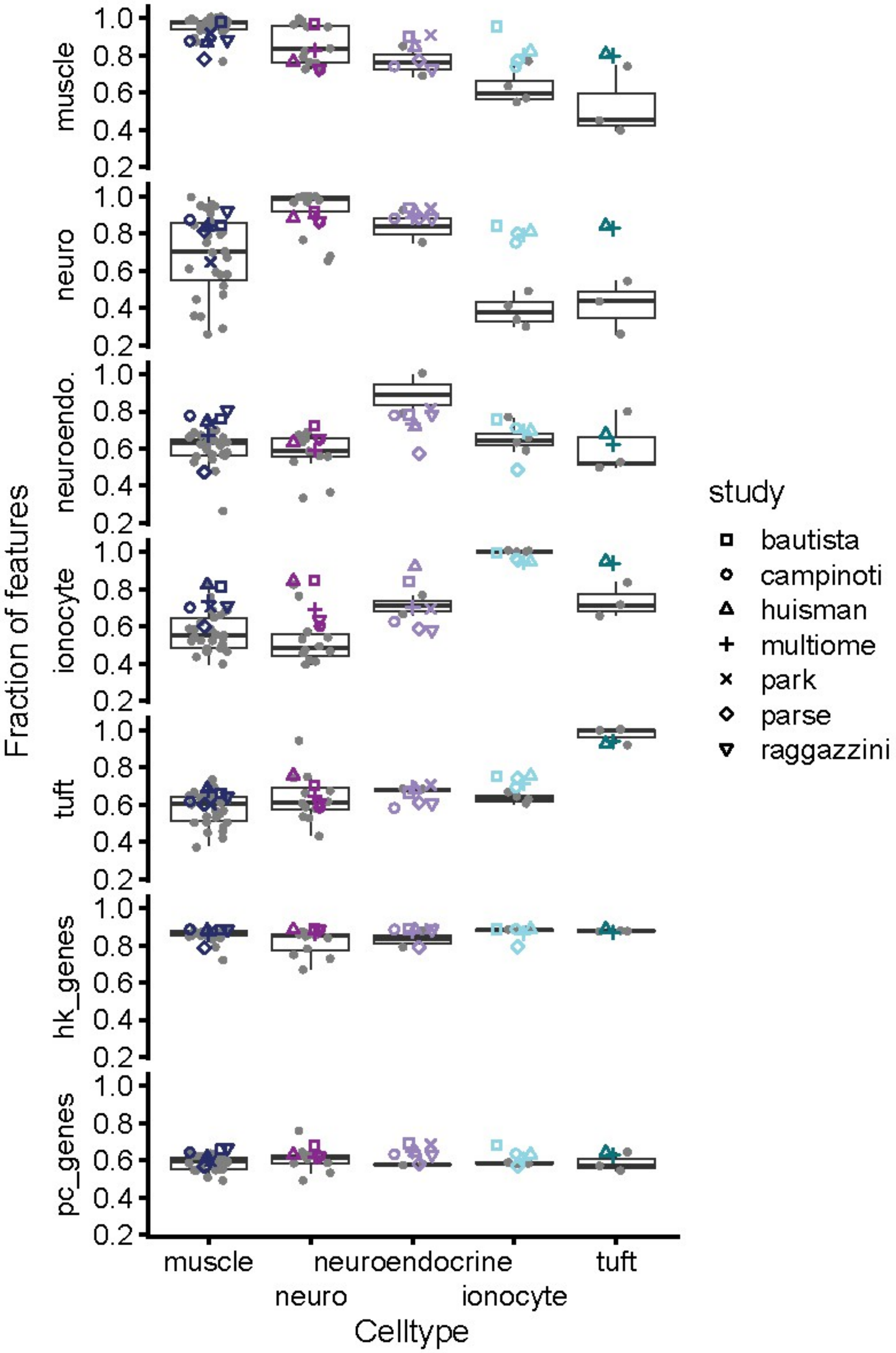


**Supplementary Figure 15.** Fraction of CRA genes expressed in TD-TECs split by study (shapes) and peripheral cell types (box and gray circles).

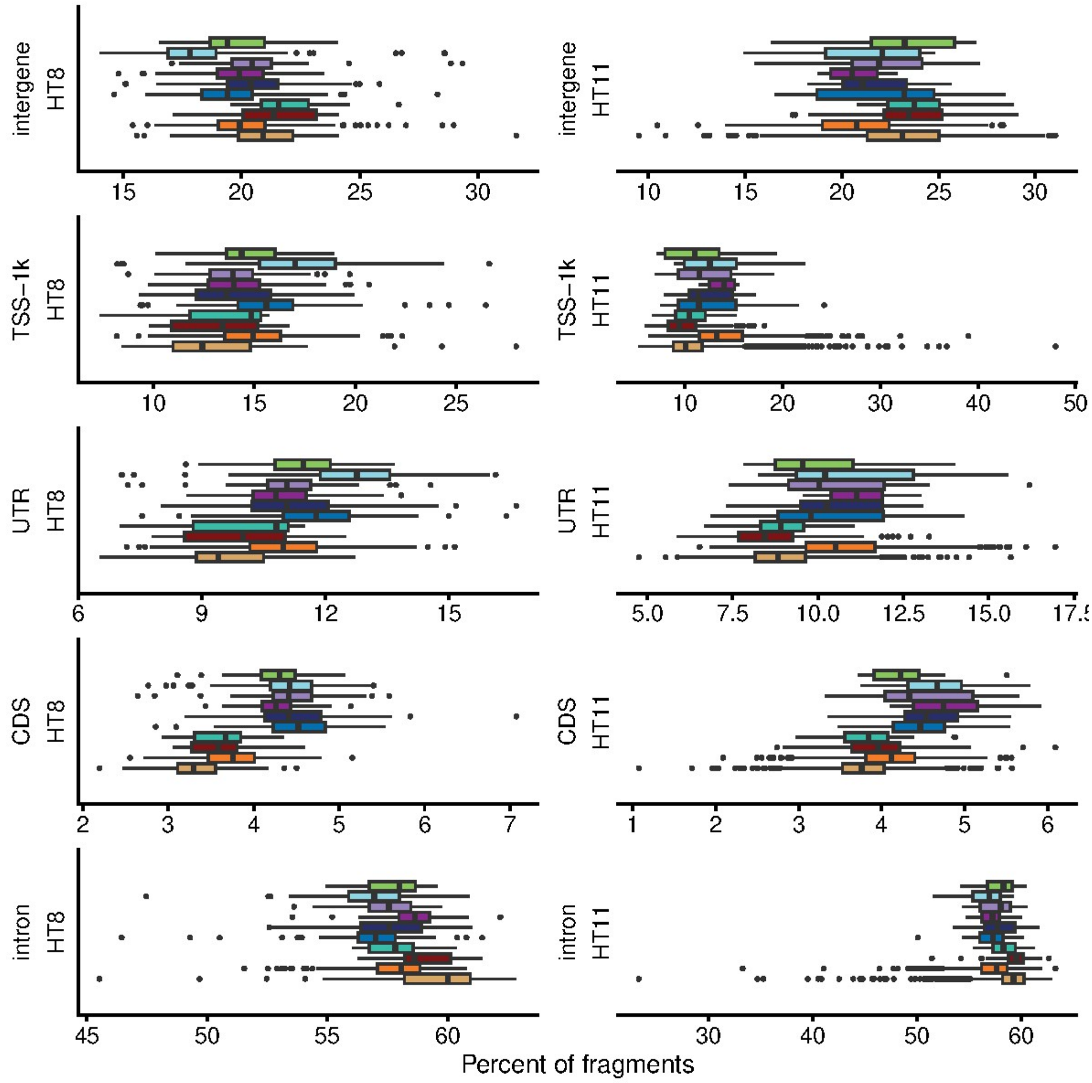


**Supplementary Figure 16.** Percent of fragments aligned to genomic regions split by sample.

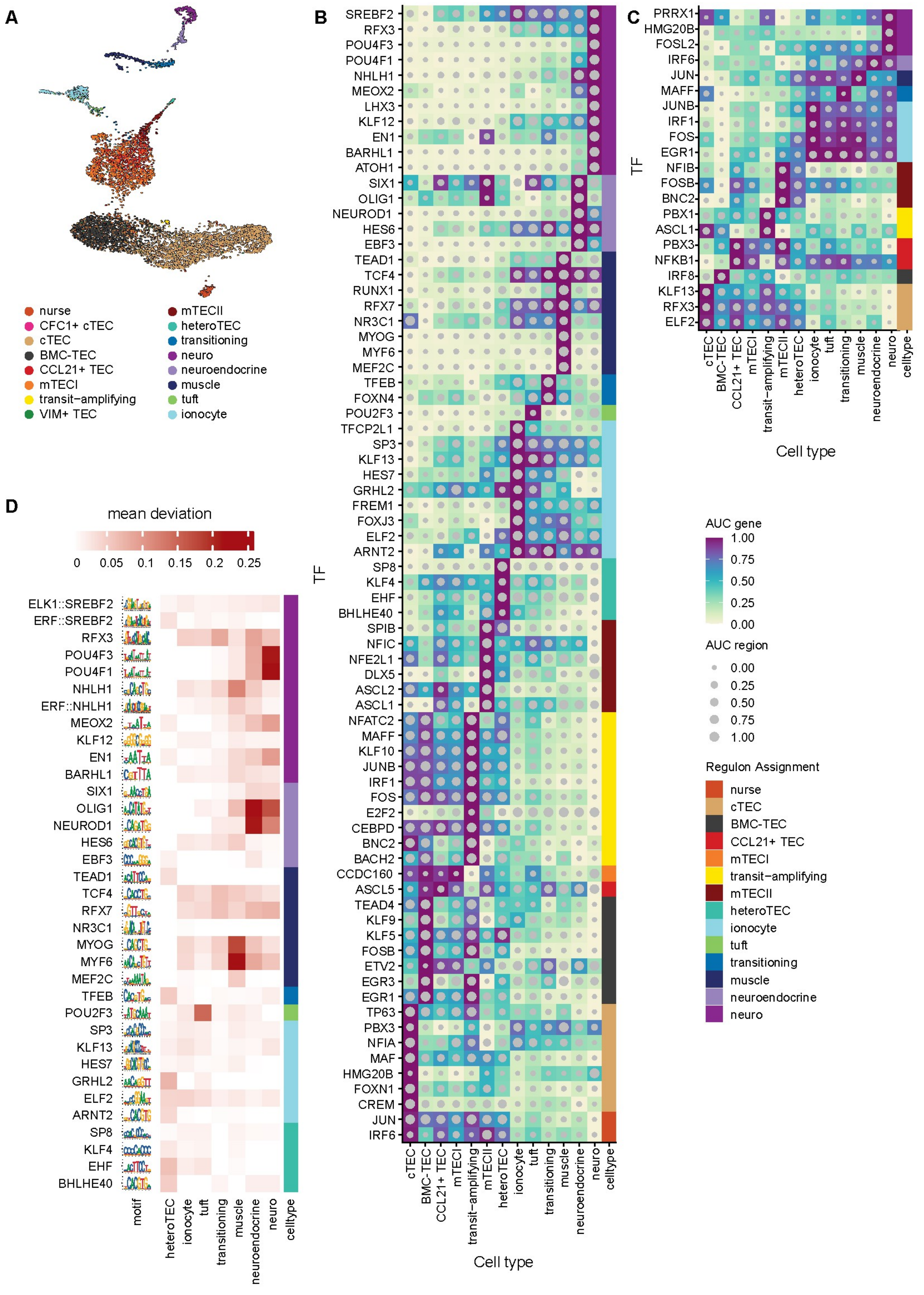


**Supplementary Figure 17. Joint analyses of accessibility and gene expression across all human TEC types. (A)** Joint UMAP embedding for paired single-nuclear RNA-seq and scATAC-seq (multiome dataset). **(A)** Joint UMAP embedding for paired single-nuclear RNA-seq and scATAC-seq (multiome dataset). **(B)** and **(C)** Combined heatmap/dotplot figures displaying gene-based and chromatin-based AUC scores for the activator (B) and repressor (C) regulons associated with all cell types. **(D)** Motif enrichment via chromVAR [13] calculated as the mean deviation score of each motif for each lineage-committed TEC type; the motifs corresponding to the activator regulon-driving TFs identified by SCENIC+ are shown.

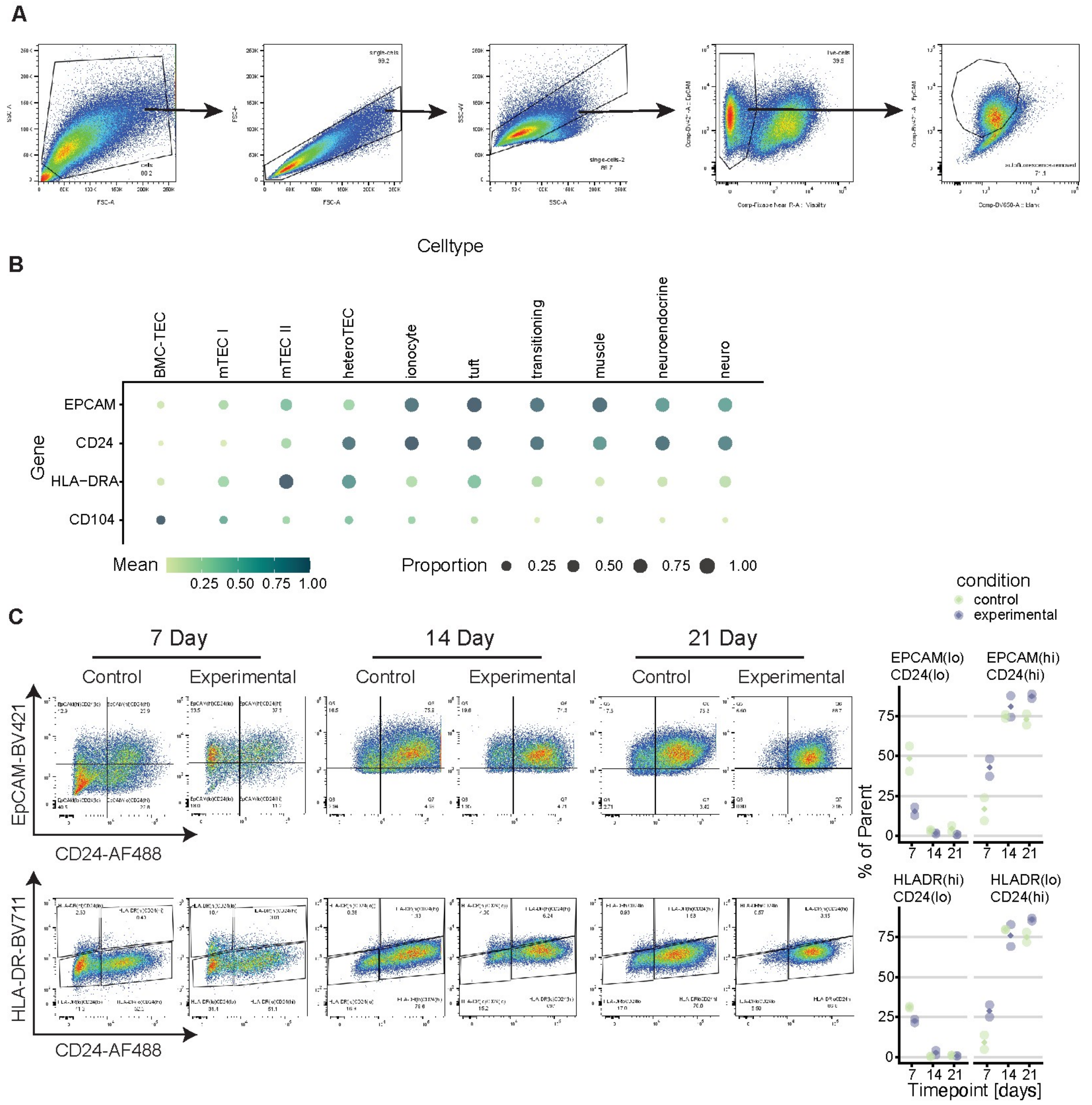


**Supplementary Figure 18.** **(A)** Sorting strategy for live sorting and culture of TECs for colony-formation and differentiation assays. **(A)** Sorting strategy for live sorting and culture of TECs for colony-formation and differentiation assays. **(B)** Bubbleplot of the expression of EPCAM, CD24, HLA-DRA, and CD104 by celltype. The color of the bubbles corresponds to the mean expression of each gene, normalized by the maximum mean expression by cell type of the given gene. The size of the bubbles corresponds to the proportion of cells in the given cell type that express the gene. **(C)** Differentiation potential of sorted HLA-DR(-/lo)CD104+CD200- TECs (BMC-TECs) after three weeks of *ex vivo* expansion in culture on an irradiated 3T3-J2 feeder layer. Expression of EpCAM, HLA-DR, and CD24 were used to assess the differentiation state of BMC-TECs at 7-, 14-, and 21-day time points under control or experimental (supplemented with retinoic acid and human RANKL) conditions. Representative flow plot (left) and quantification across biological replicates (right); diamond indicates mean.

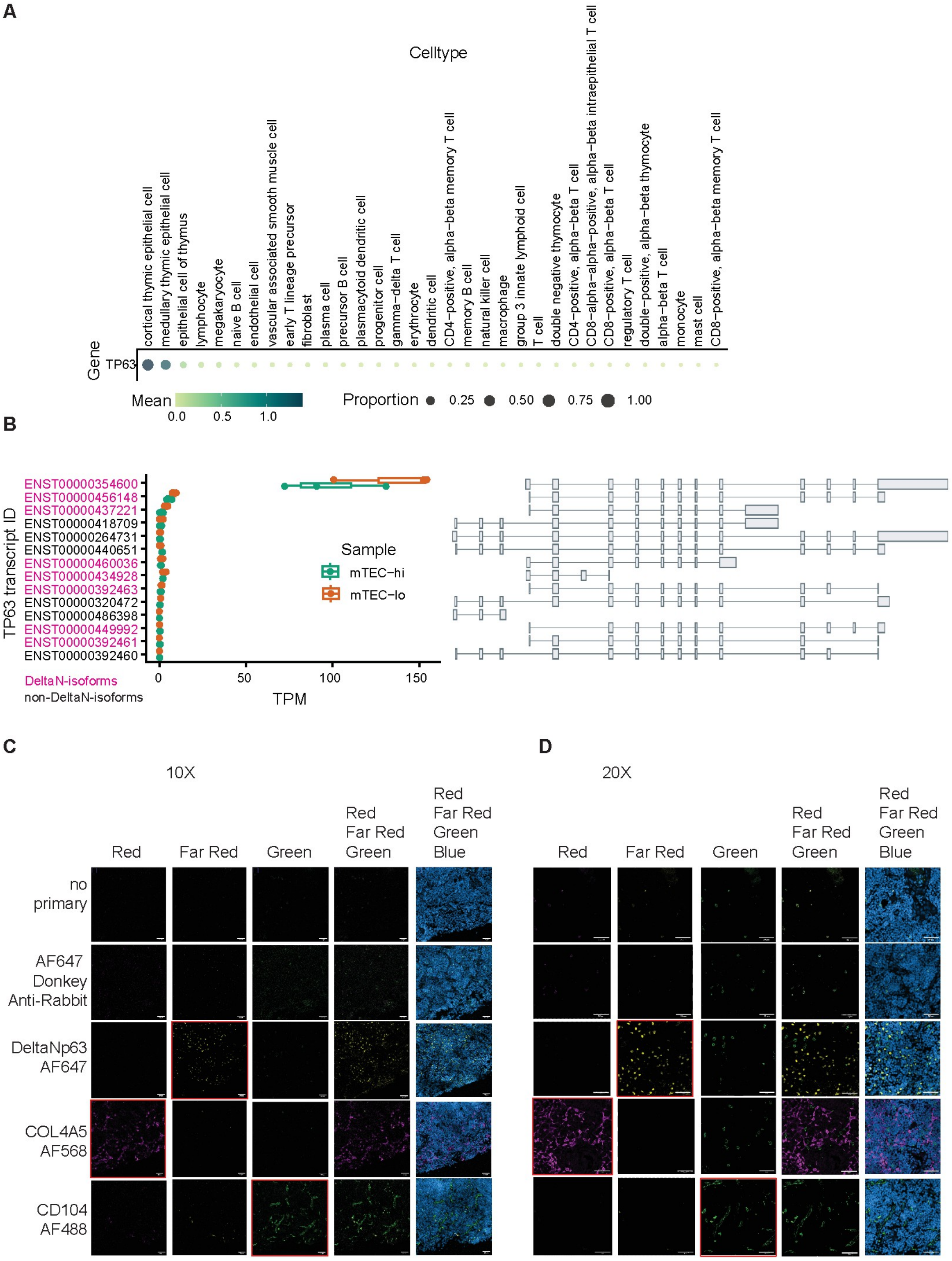


**Supplementary Figure 19. (A)** *TP63* expression in the human thymus atlas [2] (upper panel), highlighting *TP63* as TEC-specific. **(B)** *TP63* isoform expression in human mTEC [14], with DeltaN-encoding protein isoforms recognized by the rabbit anti-human DeltaN p63 antibody (Cell Signaling Technologies cat. 67825T) used for tissue staining highlighted in pink. **(C)** and **(D)** Single color controls for human thymic cryosections used for spatially locating BMC-TECs at 10x (C) and 20x (D) magnification (Scale bars: 50 µm). A 'no primary' and 'secondary only' control were used to adjust the minimum and maximum displayed pixel value. The red channel was used to detect COL4A5-AF568, the far red for DeltaNp63-AF647, the green for CD104-AF488, and the blue for DAPI; channels and corresponding fluorophores highlighted by red box. Images analyzed using Fiji version 2.16.0/1.54p.

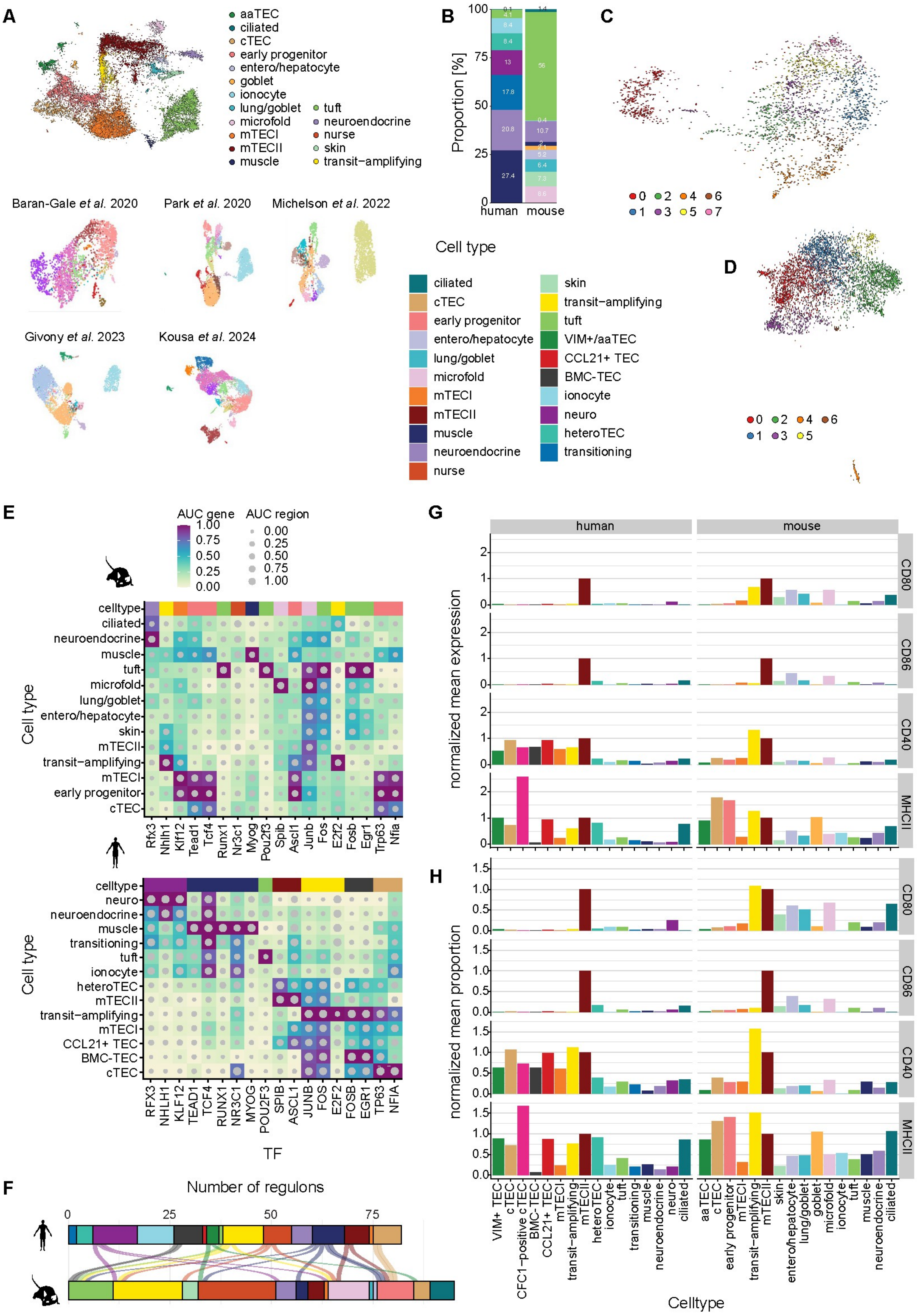


**Supplementary Figure 20. Cross-species comparison of human and mouse TD-TECs. (A)** Integrated scANVI [15] embedding of mouse TEC datasets and UMAPs of individual component datasets [2, 8, 16–18]. **(B)** Proportion of terminally differentiated TEC types from total terminally differentiated TECs in the integrated human and mouse datasets. **(C)** Integrated scVI [15] embedding of human thymus heteroTECs. **(D)** Integrated scVI [15] embedding of human thymus transitioning cells. **(E)** Combined heatmap/dotplot figures displaying mouse and human gene-based and chromatin-based AUC scores for the activator regulons conserved between the species. **(F)** Alluvial plot of all identified activator regulons and their assigned cell types in human and mouse. **(G)** and **(H)** Mean gene expression and proportion of cells per cluster expressing respective gene normalized to mean expression and proportion of AIRE/Aire-expressing mTECIIs.
